# Preclinical *CRX* augmentation therapies for *CRX*-associated autosomal dominant cone-rod dystrophies

**DOI:** 10.64898/2026.02.23.707517

**Authors:** Chi Sun, Michael Fitzpatrick, Daniel Kerschensteiner, Shiming Chen

## Abstract

Cone-rod dystrophies (CoRD) are inherited retinal diseases (IRDs) with variable ages of onset, characterized by the progressive loss of cones, followed by secondary degeneration of rods. Cone-rod homeobox (CRX) is a transcription factor that regulates gene expression essential for photoreceptor development and maintenance. Mutations in *CRX* gene, including *CRX^E168d2^* and *CRX^E80A^*, are implicated in autosomal dominant CoRDs. Although these mutations show distinct pathogenic mechanisms, published studies in knock-in mouse models have suggested a common treatment strategy: increasing WT *CRX* expression to reduce the detrimental activities of mutant proteins. This study employs two independent strategies of *CRX* augmentation to evaluate their therapeutic potential in *Crx^E168d2/+^* and *Crx^E80A/+^* mouse models. The *Tet-On-hCRX* transgenic system, a platform of proof-of-principle gene therapy, induces consistent and pan-photoreceptor expression of *CRX* augmentation in diseased retinae, allowing for the faithful assessment of functional and behavioral recovery. *AAV*-mediated *CRX* augmentation confirms the biosafety, delivery efficiency and efficacy of viral transduction in diseased retinae. Both strategies have achieved significant treatment outcomes in cone photoreceptor survival and overall photoreceptor functions in young adulthood. Treated cones survive past the age point of complete cone loss in untreated controls of both models. Treated rods show functional improvement and long-term survival through later adulthood. This preclinical study establishes valuable treatment regimens and benchmarks for *CRX* augmentation in the treatment of *CRX*-associated IRDs, and offers new insights into the mechanisms for photoreceptor development and survival.

## INTRODUCTION

Mutations in photoreceptor-specific transcription factor genes, such as the cone-rod homeobox gene (*CRX*), are associated with inherited retinal diseases (IRDs)^1^. CRX is essential for regulating the gene expression network that determines the development, function, and maintenance of vertebrate photoreceptors^2–5^. The 299-amino-acid CRX protein comprises two major functional domains, namely, N-terminal homeodomain responsible for sequence-specific DNA binding and C-terminal activation domain essential for transcriptional regulation on the target genes^2,5^. In rods, CRX forms a transcriptional complex with NRL and NR2E3 to drive the expression of rod-specific genes, such as rhodopsin (*RHO*). In cones, CRX interacts with a distinct set of transcription factors, such as RXRγ, to activate cone-specific opsin genes, *OPN1SW* and *OPN1MW*. Disease-causing mutations in the *CRX* gene produce mutant proteins with impaired DNA binding activity or transactivation ability, leading to misregulated expression of CRX-target genes. In addition, many pathogenic *CRX* mutations are inherited in the autosomal dominant pattern, causing early-onset IRDs, including amaurosis (LCA) and cone-rod dystrophy. Unfortunately, effective treatment strategies for *CRX*-associated IRDs remain an unmet medical need.

At present, gene augmentation represents the predominant therapeutic strategy for IRDs^6,7^. Gene augmentation delivers a functional copy of a gene into diseased cells to compensate for a defective endogenous allele and restore normal cellular function. In order to strategize an effective *CRX* augmentation, several key considerations must be addressed. Firstly, *CRX* augmentation must restore sufficient level of functional CRX expression to improve photoreceptor development and function in *CRX* mutants. Insufficient level of functional *CRX* expression fails to rectify the perturbed gene expression network in treated photoreceptors, causing irreversible and persistent degeneration. Secondly, *CRX* augmentation must be active during the critical treatment window in *CRX* mutants. Therapeutic intervention cannot rescue defects incurred before treatment initiation. Thirdly, overexpression of transcription factor genes may result in expression toxicity, disrupting the precisely regulated gene expression network. Lastly, *CRX* expression is indispensable to developing and mature photoreceptors. An effective *CRX* augmentation must maintain robust and long-term expression. So far, *CRX* augmentation has been assessed in human retinal organoid models of *CRX*-associated LCA^8^. However, whether *CRX* augmentation can effectively promote photoreceptor function and survival remains unclear.

This study determines the therapeutic potential and preclinical benchmarks of *CRX* augmentation by two independent systems in distinct models of early-onset *CRX*-associated cone-rod dystrophies. *Tet-On-hCRX* is a transgene system designed to induce a pan-photoreceptor and consistent *CRX* expression, facilitating a faithful assessment of the treatment efficacy and functional recovery in treated mice. *AAV*-mediated CRX augmentation is employed to evaluate the therapeutic dose range and optimal promoters for *CRX* expression. Results of this study provide critical insights for translating effective therapeutics targeting diseased photoreceptors of *CRX*-associated IRDs.

## METHODS

### Transgenic mouse lines

Mice were on the genetic strain of *C57BL/6J. Tet-On-hCRX* consisted of three components. (1) *TRE-hCRX* transgene contained a *TRE* promoter upstream of full-length human *CRX* cDNA sequence. The transgene was inserted into the *H11* locus. (2) *pCAG-LSL-rtTA* (also known as *Rosa26:pCAG-LSL-rtTA*) was obtained from the Jackson Laboratory (RRID:IMSR_JAX:029617). (3) *pCrx-Cre* (also known as BAC-Tg *Crx-Cre*) was a published colony^9^. Doxycycline-containing chows (200mg/kg, Bio-Serv S3888) were provided to *Tet-On-hCRX* mice including breeding dams. Augmented *hCRX* expression can be detected at neonatal periods, indicating dox administration in pups through milk or other contacts^10,11^. 100μL of 20mg/mL doxycycline solution was intraperitoneally injected into *Tet-On-hCRX* mice (>P21) once a week to maximize the treatment effect.

*Tet-On-hCRX* functions in following way. *pCrx-Cre* induces Cre recombinase in both classes of photoreceptors. Cre recombinase activates *rtTA* expression in *pCAG-LSL-rtTA*. Doxycycline binds to trigger a conformational change to the rtTA protein, allowing the complex to bind to *TRE* element in *TRE-hCRX*. Ultimately, *hCRX* expression is activated. Human CRX protein is tagged with 3XFLAG. A schematic diagram is illustrated in Supplementary Figure 19. DNA construct of *TRE-hCRX* is provided in Supplemental table 1. Genotyping primers for *TRE-hCRX* transgene are F (5’ to 3’) CACAGGTGTCCACTCCCAGTTCAATTAC, R (5’ to 3’) CTTAGGGCCAAGGCGTTGACAGAATAG.

*Crx^E168d2/+^* and *Crx^E80A/+^* mice were generated in published studies^12,13^. Bot male and female mice were used in the experiments.

### AAV production and delivery

Four types of *AAV* constructs were used in this study: *AAV*-*pGnb3*-Tet-Off-hCRX*, *AAV*-*pGnb3*-Tet-Off-DsRed*, *AAV*-*pGrk1-Tet-Off-hCRX*, *AAV*-*pGrk1-Tet-Off-DsRed*. Details of *AAV* constructs were included in Supplemental table 2. *AAV* constructs were packed into *AAV2/5* vectors at Hope Center Viral Vectors Core, Washington University in St. Louis.

Mice were anesthetized with xylazine (10 mg/kg) and ketamine (100 mg/kg) saline solution prior to subretinal injections. Subretinal injection was performed under a stereomicroscope (Leica). A limbal hole was firstly made by a 31G needle, viral solutions were subsequently delivered to the subretinal space through the hole by a blunt 33G Hamilton syringe. Viral solutions had type concentrations of 10^13^ vg/mL. Each eye of a mouse was injected up to 0.5 μL of viral solutions. Successful injections should show no significant vitreous or subretinal hemorrhage.

### Treatment schemes

A summary of *Tet-On-hCRX* and *AAV-hCRX* treatments is illustrated in Supplemental figure 20.

### Quantitative PCR (qPCR)

Each RNA sample was extracted from 2 retinae of a mouse using the NucleoSpin RNA Plus kit (Macherey-Nagel). 2 μg of RNA was used to produce cDNA using First Strand cDNA Synthesis kit (Roche). qPCR Primers were listed in Supplemental table 3. The reaction master mix contained EvaGreen polymerase (Bio-Rad Laboratories), 1 μM primer mix, and diluted cDNA samples. qPCR was done with a two-step 40-cycle protocol on a Bio-Rad CFX96 Thermal Cycler (Bio-Rad Laboratories). Pairwise t-test was performed with p < 0.05, CI:95% using Graphpad Prism 10 (GraphPad Software).

### Western Blotting

Protein lysates were obtained using NE-PER™ Nuclear and Cytoplasmic Extraction Reagents (ThermoFisher Scientific). Protein concentrations were determined using Pierce™ BCA Protein Assay Kit (ThermoFisher Scientific) on a NanoDrop One spectrophotometer (ThermoFisher Scientific). Either 10ug of nuclear protein lysate or 20ug of cytoplasmic protein lysate was used for Western blotting. Protein samples were boiled in water for 10min and then run in NuPAGE™ Bis-Tris 4–12% Gels (Invitrogen). Samples were transferred onto PVDF membranes using Mini Blot module (Invitrogen). Membranes were blotted using iBind system (Invitrogen). Western blotting of anti-CRX (1:500), anti-FLAG (1:500), anti-Lamin B1 (1:1000) was done with nuclear protein lysates; Western blotting of anti-ARR3 (1:300) was done with cytoplasmic protein lysates. Information on primary antibodies was included in Supplemental table 4. Goat anti-mouse IRDye and anti-rabbit IRDye (LI-COR) were selected as secondary antibodies. Imaging was done with Odyssey CLx Imaging system (LI-COR).

### H&E and Immunohistochemistry (IHC) Staining

Dissected eyes were fixed in 4% paraformaldehyde for 1 or 1.5 hour, and then prepared for paraffin-embedded or cryo sections. Paraffin-embedded sections were used in H&E staining and IHC staining of RXRγ, RHO, and GFAP. Cyro sections were used in IHC staining of ARR3 and CC3.

For preparing cryo sections, fixed tissues were incubated in increasing concentrations (5% to 20%) of phosphate-buffered sucrose solutions, and then frozen in OCT compound (optimal cutting temperature, ThermoFisher Scientific). Cyro sections were cut by 10 microns. Paraffin-embedding was prepared in Sakura Tissue-Tek VIP tissue processor. H&E staining was performed with 4-micron paraffin-embedded sections. Unstained paraffin-embedded sections were incubated with hot citrate buffer for antigen retrieval prior to IHC staining procedures.

IHC staining followed the protocol of the previous study^14^. Blocking buffer contained 5% donkey serum and 0.1% Triton X-100 in 1X phosphate-buffered saline (PBS) (pH 7.4). Slides were incubated with selected primary antibodies at 4°C overnight. Slides were washed with 1X PBS for 30 min, and then incubated with specific secondary antibodies for 2 hours. Slides were washed and mounted with hard set mounting medium with DAPI (Vectashield, Vector Laboratories). Primary antibodies are listed in Supplemental table 4.

All images were taken on a Leica DB5500 microscope. All images were acquired at 1000-1500μm from optical nerve head for ≥ P14 samples and at 500μm from optical nerve head for < P14 samples. Thickness and cell count analysis was performed as follows. ONL and OS thickness was measured at defined distance from the optical nerve head on both the superior and inferior retina. Mean values of measurements on biological replicates were plotted. Pairwise t-test or ANOVA weas performed with performed with p < 0.05, CI:95% using Graphpad Prism 10 (GraphPad Software). Number of fluorescent cells were counted in defined regions as described in figure legends. Pairwise t-test was performed with p < 0.05, CI:95% using Graphpad Prism 10 (GraphPad Software).

### Electroretinography (ERG)

Mice were dark-adapted overnight prior to ERG tests. ERG followed the protocol of the previous study^14^. ERG tests were performed with UTAS-E3000 Visual Electrodiagnostic System (LKC Technologies). ERG protocol was described in the previous study^14^. In brief, dark-adapted responses were recorded with full-field light flashes (10 μs) of increasing intensity (totally 9 intensities), the maximum flash intensity was 0.895 cd*s/m^2^. Light-adapted responses were measured with 7 intensities, maximum flash intensity for light-adapted testing was 2.672 cd*s/m^2^. ERG responses of biological replicates were recorded, averaged, and analyzed using GraphPad Prism 10 (GraphPad Software). The mean peak amplitudes were plotted against log values of light intensities (cd*s/m^2^). The statistics were analyzed by ANOVA or pairwise t-test.

### Optokinetic reflex (OKR) and pupillary light reflex (PLR)

OKR evaluates the reflexive ability of the eye to track drifting gratings, quantified as the number of eye-tracking movements (ETMs) per 60 seconds of the stimuli. PLR measures constriction of the pupil to steps of varying light intensity. The transient phase of the PLR requires photoreceptor input^15,16^. OKR and PLR methodology is illustrated in Supplementary Figure 21. In this study, OKR and PLR tests follow the published protocol and procedures^17^. For OKR experiments, visual stimuli were presented using the Cogent Graphics toolbox in MATLAB (The MathWorks). Square wave gratings of varying spatial frequencies (0.05, 0.067, 0.1, 0.13, 0.2 cpd) moving at 10°/s in the temporal-to-nasal direction were presented on a monitor 16 cm from the mouse’s left eye at a 45° angle. Each stimulus of OKR presentation contained 10 seconds of a uniform gray screen, 60 seconds of drifting gratings, and a final 10 seconds of a uniform gray screen. Eye-tracking movements were quantified as the number of saccades preceded by a slow tracking motion. Each stimulus of PLR presentation comprised 5 seconds of background darkness, 30 seconds of illumination, and 30 seconds of post-illumination recovery to baseline, with a minimum of 2 minutes in darkness between presentations. Pupil constriction was normalized to the dark-adapted pupil area and calculated as a 5 second average around the point of maximal constriction. EC50 values were derived by fitting a Hill equation to the constriction at each illuminance for each animal.

## RESULTS

### Photoreceptor degeneration in mouse models for *CRX*-associated cone-rod dystrophies

Cone-rod dystrophy manifests with early-age cone defects, such as reduced visual acuity and frequent dyschromatopsia, followed by concomitant rod degeneration^18–20^. Considering the syndromic severity, a feasible treatment strategy for cone-rod dystrophy has wide application in treating other forms of IRDs. Since CRX is expressed in both classes of photoreceptors, *CRX*-associated cone-rod dystrophy serves as an ideal disease model for evaluating treatment strategies targeting cones and rods. Both *CRX^E168d2/+^* and *CRX^E80A/+^* are linked to early-onset autosomal dominant cone-rod dystrophy^21,22^. These two mutations represent distinct classes of *CRX* mutations and give rise to different disease phenotypes. Therefore, it is important to determine the detailed courses of photoreceptor degeneration in knock-in mouse models of *Crx^E168d2/+^* and *Crx^E80A/+^* to establish untreated baselines for therapeutic evaluation.

The frameshift *CRX^E168d2^* mutation is located within the coding region for the activation domain, producing a truncated protein, p.E168d2, that retains DNA binding ability but fails to regulate target gene expression^12^. The mouse model of *Crx^E168d2/+^* shows retinal phenotypes similar to the key clinical manifestations observed in human patients^12^. Hematoxylin and eosin (H&E) staining of retinal sections reveals progressive retinal degeneration in adult *Crx^E168d2/+^* mice (Supplemental figure 1A). Notably, the structure of outer segments (OS) is almost entirely lost in 5 month-old (MO) *Crx^E168d2/+^* retinae. As compared to wildtype (*WT*) controls, mutant retinae show a significant decrease in outer nuclear layer (ONL) thickness beginning at 3MO, suggesting the extensive photoreceptor degeneration in early adulthood (Supplemental figure 1B). Cone degeneration is not evident before postnatal day (P) 14 in *Crx^E168d2/+^* retinae (Supplemental figure 1C, 1E) but rapidly advances following the completion of photoreceptor differentiation (Supplemental figure 1D, 1E). As compared to *WT* controls, *Crx^E168d2/+^* retinae retain only 34% survived cones at 1 MO and no detectable cones by 3MO. It has been shown that p.E168d2 is overproduced relative to the WT CRX protein in *Crx^E168d2/+^* retinae^12^. The corresponding *Crx* transcript expression follows a similar pattern. In *Crx^E168d2/+^* retinae, total *Crx* transcript expression (i.e. *Crx^E168d2^* and *WT Crx* transcripts) gradually increases from 1MO (Supplemental figure 1F). In contrast, *WT Crx* transcript expression becomes downregulated from 1MO (Supplemental figure 1G), resulting in a declined ratio of *WT*:*Crx^E168d2^ Crx* gene expression. Given that ONL thickness remains comparable in *Crx^E168d2/+^* retinae at these tested ages, this imbalanced ratio likely results from differential expression in *Crx^E168d2^* and *WT Crx* transcripts rather than from altered cell numbers.

The substitution *CRX^E80A^* mutation lies within the coding region for the homeodomain, resulting in a mutant protein, p.E80A, with altered DNA binding capacity^13^. Unlike *Crx^E168d2/+^* retinae, *Crx^E80A/+^* retinae do not exhibit a rapid reduction in ONL thickness in early adulthood (Supplemental figure 2A, 2B). However, cone defects occur soon after P0^13^. Retinoid X receptor gamma (RXRγ) is expressed in developing cones of neonatal *WT* retinae. Anti-RXRγ Immunohistochemistry (IHC) identifies that *Crx^E80A/+^* retinae contain only 54% RXRγ+ cones as compared to *WT* controls at P3, suggesting the critical treatment window for early-stage cone differentiation (Supplemental figure 2C). While differentiating cones express cone arrestin (ARR3) in *WT* controls at P10, only a few ARR3+ cones can be detected in *Crx^E80A/+^* retinae. Cone populations undergo a rapid loss during cone differentiation, with no cones detectable at P21 (Supplemental figure 2C, 2D). Molecular analysis has shown that p.E80A shares the same DNA binding specificity with the WT CRX protein^13^ but fails to dynamically facilitate the transcription activity at dimeric K50 motifs that drive cone-enriched gene expression^23^. Despite their different pathogenic mechanisms, both *CRX^E168d2^* and *CRX^E80A^* mutations cause early-onset cone loss followed by later rod degeneration. This pattern of disease progression hence highlights the need for early and sustained therapeutic interventions.

### Development of the *Tet-On-hCRX* transgenic system

*CRX* augmentation can be a viable treatment strategy for *CRX^E168d2/+^* and *CRX^E80A/+^* mutants. Photoreceptor phenotypes are consistently less severe in heterozygous mouse models of these mutations as compared to homozygous ones^12,13^. In particular, reduced expression of *Crx^E168d2^* allele correlates with a less severe phenotype and a delayed age of onset^12^. More importantly, a primary pathogenic mechanism in heterozygous mouse models involves insufficient levels of functional WT CRX protein. *CRX* augmentation directly addresses this deficit. Augmented *CRX* expression may improve the transcription activation at CRX-target genes, thereby counteracting the dominant-negative effect by the nonfunctional mutant protein in *Crx^E168d2/+^* retinae. On the other hand, augmented *CRX* expression may enhance the binding activity of p.WT CRX protein at dimeric K50 motifs to promote cone differentiation in *Crx^E80A/+^* retinae. Therefore, the running hypothesis of this study states that CRX augmentation can promote the transcription activation at CRX-target genes, thereby counteracting the dominant-negative effect by nonfunctional mutant CRX proteins.

This study aims to achieve two main objectives to assess the efficacy of *CRX* augmentation. Considering the earlier onset of cone degeneration, gene therapy strategies for *CRX*-associated cone-rod dystrophies should prioritize cone survival. An optimal gene therapy strategy should also improve rod function and survival, thereby generating a neuroprotective microenvironment for overall photoreceptor viability.

A transgenic mouse line, *Tet-On-hCRX*, has been generated to activate inducible *CRX* augmentation in photoerceptors^24^. A detailed schematic of this transgenic system is illustrated in Methods. In brief, the *Tet-On-hCRX* system contains three components: *pCrx-Cre*, *pCAG-LSL-rtTA*, *TRE-hCRX*. In the presence of doxycycline (dox), reverse tetracycline-controlled transcriptional activator (rtTA) binds the synthetic *tetracycline-responsive element* (*TRE*)^25,26^, activating human *CRX* (*hCRX*) gene expression (Figure 1A). *CrxCre*^9^ ensures *hCRX* expression in photoreceptors. Augmented CRX protein is FLAG-tagged, allowing for uniform detection by anti-FLAG IHC in nuclei of *Tet-On-hCRX*-treated *Crx^E168d2/+^* photoreceptors (denoted as *Crx^E168d2/+^*^,*Tet-On-hCRX*^ in following sections, Figure 1B). These results altogether confirm the inducibility of the *Tet-On-hCRX* system even in diseased photoreceptors.

**Figure 1.**
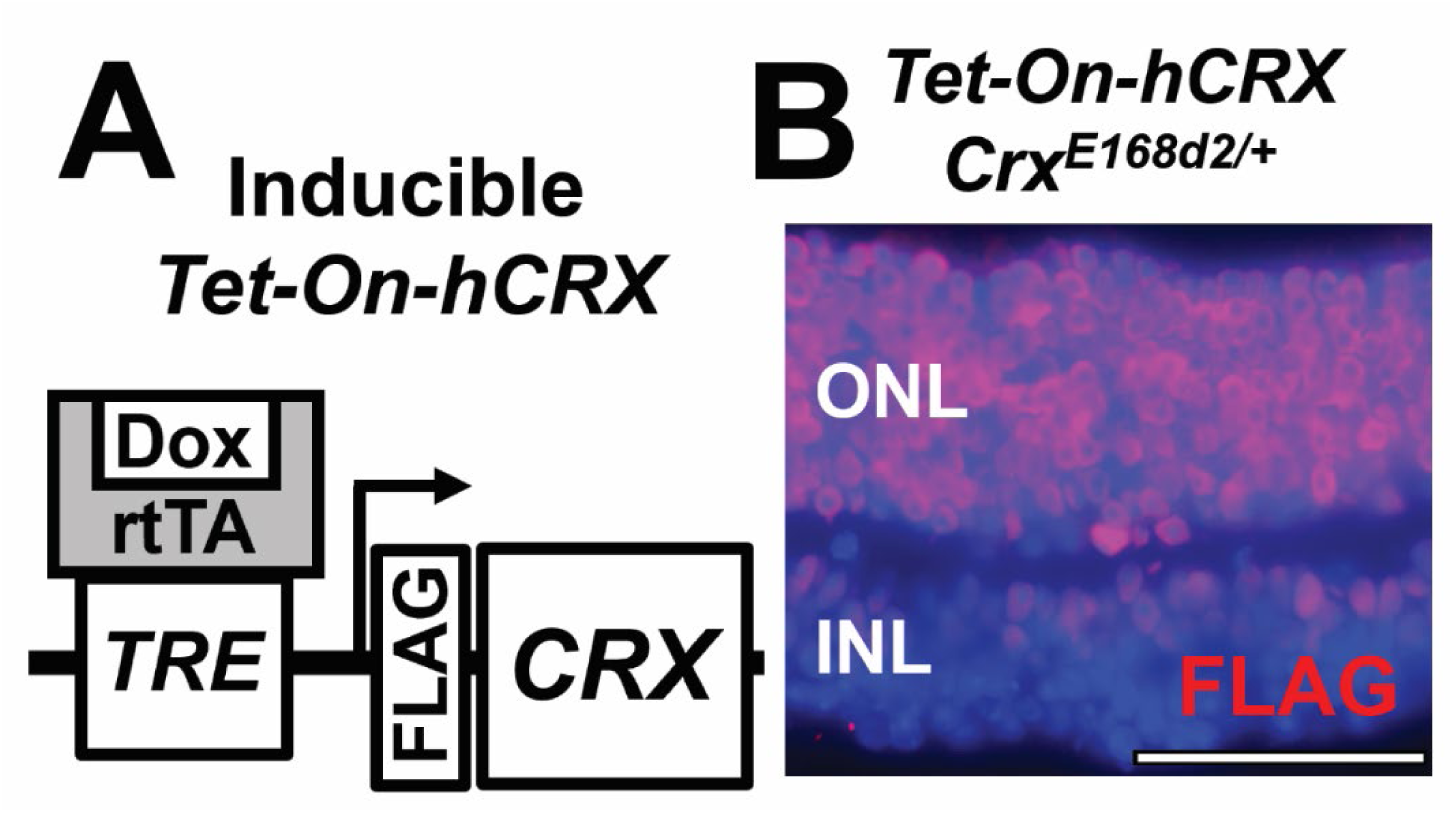
*Tet-On-hCRX*. (A) The *Tet-On* system for doxycycline-inducible human *CRX* (*hCRX*) expression. Augmented CRX is tagged with FLAG. (B) Anti-FLAG IHC staining (in red) in *Crx^E168d2/+^*^;*Tet-On-hCRX*^ retinae at 1MO. Nuclei are stained by DAPI (in blue). Scale bar = 50µm. ONL: outer nuclear layer. INL: inner nuclear layer.

It is important to validate the safety of *Tet-On-hCRX* system prior to evaluating its therapeutic potential. Augmented CRX protein is detectable in *WT^Tet-On-hCRX^* retinae at P5 (data not shown). 1MO *WT^Tet-On-hCRX^* retinae exhibit normal retinal morphology (Supplemental figure 3A), whose ONL and OS thickness (Supplemental figure 3B) is comparable to that of *WT* controls. Furthermore, *Tet-On-hCRX* system shows no adverse effects on electroretinogram (ERG) responses (Supplemental figure 3C), suggesting good tolerability of *CRX* augmentation in treated photoreceptors.

### *Tet-On-hCRX* improving *Crx^E168d2/+^* photoreceptor morphology, function and survival

The *Tet-On-hCRX* system is employed to treat *Crx^E168d2/+^* retinae. Doxycycline can be administrated at different ages. In order to achieve early treatment, pups receive doxycycline via placenta (embryonic period) and lactation (neonatal period) from treated nursing dams. *CRX* augmentation by *Tet-On-hCRX* can be detectable by Western blotting on FLAG in developing *Crx^E168d2/+^*^,*Tet-On-hCRX*^ retinae at P5 (Supplemental figure 6A). Developing RXRγ+ cones are localized to apical regions of P0 *Crx^E168d2/+^*^,*Tet-On-hCRX*^ retinae (Supplemental figure 4A), showing comparable cell localization and numbers to those in *WT* and *Crx^E168d2/+^* retinae (Supplemental figure 4B). This suggests that *Tet-On-hCRX* system does not alter the intrinsic development program in early cones. At 3MO, *Crx^E168d2/+^*^,*Tet-On-hCRX*^ retinae exhibit well-organized morphology (Figure 2A) and significant increase in ONL and OS thickness as compared to untreated retinae (Supplemental figure 5A, 5B). Anti-RHO IHC confirms longer rod OS in *Crx^E168d2/+^*^,*Tet-On-hCRX*^ retinae (Figure 2B), while anti-ARR3 IHC detects a population of survived cones in treated retinae (Figure 2C) in contrast to complete cone loss in untreated retinae (Supplemental figure 5C). These results demonstrate that *CRX* augmentation via *Tet-On-hCRX* significantly improves retinal morphology and cone survival in treated *Crx^E168d2/+^* retinae.

**Figure 2.**
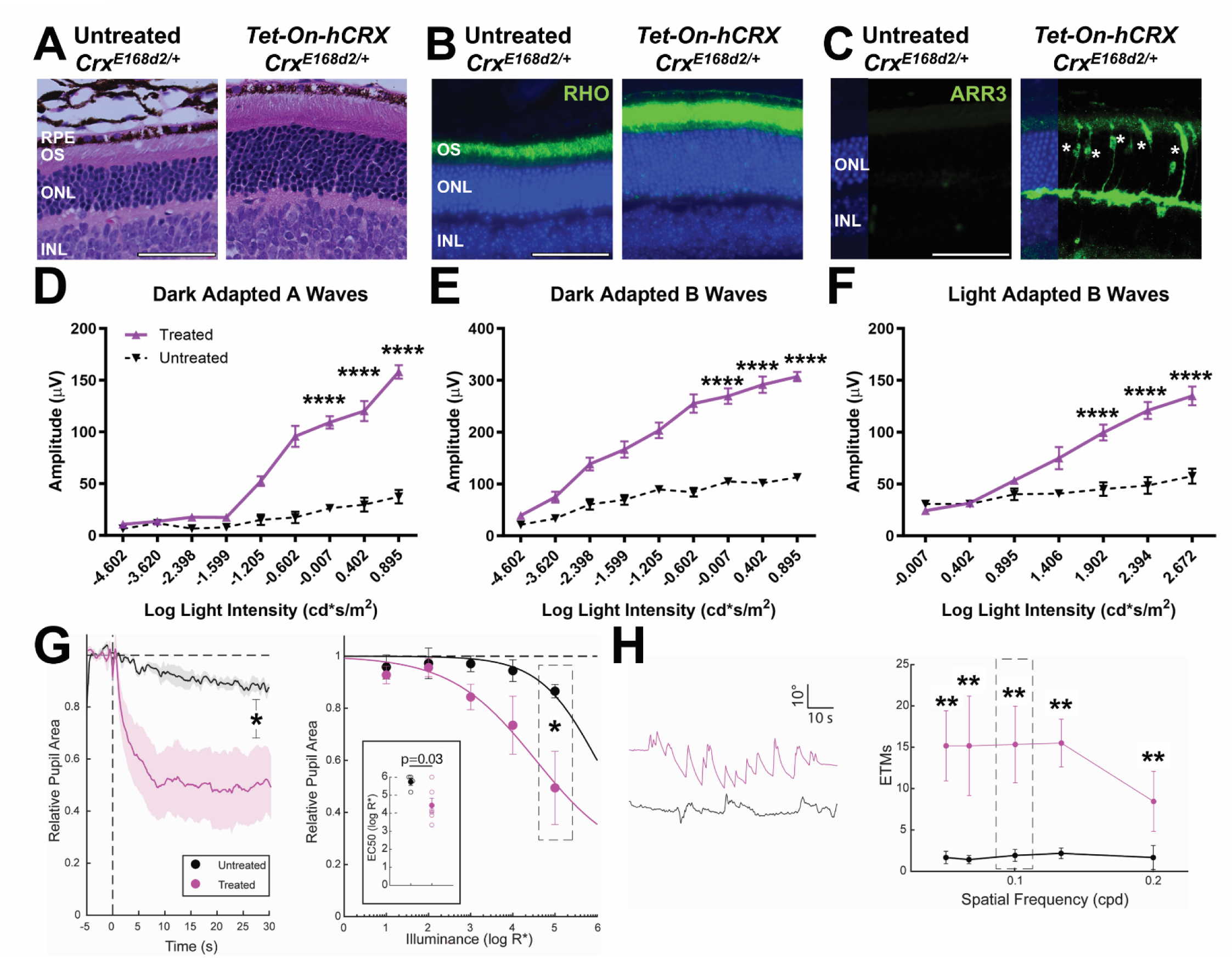
*Tet-On-hCRX*-mediated *CRX* augmentation rescuing *Crx^E168d2/+^* retinae. (A) H&E cross-section staining of *Crx^E168d2/+^* and *Crx^E168d2/+;Tet-On-hCRX^* retinae at 3MO. RPE: retinal pigment epithelium. OS: outer segment. Scale bar = 50µm. (B) Anti-RHO IHC staining (in green) in *Crx^E168d2/+^* and *Crx^E168d2/+;Tet-On-hCRX^* retinae at 3MO. Nuclei are stained by DAPI (in blue). (C) Anti-ARR3 IHC staining (in green) in *Crx^E168d2/+^* and *Crx^E168d2/+;Tet-On-hCRX^* retinae at 3MO. Asterisks indicate ARR3+ cones in *Crx^E168d2/+;Tet-On-hCRX^* retinae. (D) ERG measurements of dark-adapted A waves of *Crx^E168d2/+^* and *Crx^E168d2/+;Tet-On-hCRX^* mice at 3MO. Error bars represent SD on mean values (n=8). (E) ERG measurements of dark-adapted B waves. (F) ERG measurements of light-adapted B waves. (G) Pupillary light reflex (PLR) measurements. The left panel details average pupil constriction for treated vs untreated animals at the brightest intensity tested, as marked by the dotted box in the overall-mean plot (right). Error bars represent SEM (n=6). (H) Optokinetic response (OKR) measurements. The left panel details representative plots for a treated vs an untreated animal at 0.1 cpd, showing an enlarged view of the region marked by the dotted box in the overall-mean plot (right). Error bars represent SEM (n=8). Statistical analysis by ANOVA pairwise t-test happens between *Crx^E168d2/+^* and *Crx^E168d2/+;Tet-On-hCRX^* samples. Statistical significance in this figure is indicated by asterisks (*, **, ****) denoting p≤0.05, p≤0.01, and p≤0.0001 respectively, and ns meaning not significant.

It is critical to assess if survived photoreceptors recover functional capacity. ERG directly measures the electrical response of treated photoreceptors, providing evidence that these cells restore visual functions beyond structural improvements. Treated photoreceptors show significantly improved rod-driven ERG (Figure 2D,E) and cone-driven ERG (Figure 2F) as compared to untreated ones. Pupillary light reflex (PLR)^17^ and acuity-related optokinetic reflex (OKR)^27^ are employed to evaluate visual behaviors in treated animals. PLR measures the pupillary constriction in response to light stimuli, assessing the integrity of photoreceptors and downstream neural pathways. OKR measures the ability to track moving stimuli, monitoring visual acuity and motion perception. *Crx^E168d2/+^*^,*Tet-On-hCRX*^ retinae display significant improvements in PLR (Figure 2G) and OKR (Figure 2H) as compared to untreated retinae. These results indicate that treated photoreceptors remain functionally integrated within the visual circuit, showing enhanced input-output responses in signal transmission and visual-motor learning.

Moreover, the *Tet-On-hCRX* system reduces glial activation in *Crx^E168d2/+^*^,*Tet-On-hCRX*^ retinae prior to the extensive photoreceptor degeneration (Supplemental figure 7A). Therefore, the amelioration of cellular and molecular pathology in *Crx^E168d2/+^*^,*Tet-On-hCRX*^ retinae leads to a corresponding decrease in apoptosis-related cell death (Supplemental figure 7B).

Sustained *CRX* augmentation is required for the effective treatment of *Crx^E168d2/+^* retinae. Doxycycline administration is withdrawn from *Crx^E168d2/+^*^,*Tet-On-hCRX*^ at 1MO (Supplemental figure 8A), thereby inactivating the *Tet-Off-hCRX* system within the onset of cone degeneration. ARR3+ cones are still present in samples 1 month post-withdrawal (Supplemental figure 8B), showing more abundance than untreated retinae. These results suggest a critical treatment window before 1MO to achieve long-lasting cone survival. However, following the loss of CRX augmentation, diseased cones undergo accelerated degeneration as compared to continuously treated samples, causing a more significant drop in cone populations by 1 month post-withdrawal (Supplemental figure 8B, 8C).

The efficacy of CRX augmentation follows ‘earlier-the-better’ principle in *Crx^E168d2/+^* retinae. The comparison involves two starting ages: early treatment (initiated at birth) and late treatment (initiated at P21). Late treatment begins at P21 when diseased cones undergo early-stage degeneration (Supplemental figure 1E), mimicking the conditions of early-onset syndromes in patients. At 3MO, early treatment results in significantly improved retinal morphology in terms of ONL and OS thickness as compared to late treatment (Supplemental figure 9A, 9B, 9C). Cone survival is also promoted by late treatment (Supplemental figure 9E), albeit less effectively than by early treatment (Supplemental figure 9F). Nonetheless, both early and late treatments yield functional recovery, as measured by improved ERG responses in treated retinae. (Supplemental figure 9G). The superior functional recovery with early treatment well agrees with its enhanced morphological assessments.

*CRX* augmentation does not halt the photoreceptor degeneration but effectively delays its progression. At 1MO, *Crx^E168d2/+^*^,*Tet-On-hCRX*^ retinae display well-preserved ONL although with slightly shorter OS as compared to *WT* controls (Supplemental figure 10A, 10G). At 3MO and 5MO, *Crx^E168d2/+^*^,*Tet-On-hCRX*^ retinae show a progressive loss of ONL and OS thickness, but retain >50% of the thickness as compared to *WT* controls. (Supplemental figure 10C, 10E, 10G). In contrast to severely lost OS in 5MO *Crx^E168d2/+^* retinae (Supplemental figure 1A), the morphology of *Crx^E168d2/+^*^,*Tet-On-hCRX*^ retinae highlights the long-term photoreceptor rescue by the *Tet-Off-hCRX* system. Long-term photoreceptor rescue is also evident by survived cones in 5MO *Crx^E168d2/+^*^,*Tet-On-hCRX*^ retinae by 5MO (Supplemental figure 10B, 10D, 10F), as compared to complete cone loss in 3MO *Crx^E168d2/+^* retinae (Supplemental figure 1D). However, the slowed but progressive degeneration leads to a drop in cone populations reduced to 78%, 41%, and 24% of *WT* controls at 1MO, 3MO, and 5MO respectively (Supplemental figure 10H).

### *CRX* augmentation promoting CRX-target gene expression

Western blotting confirms the consistent expression of FLAG-tagged CRX and ARR3 in *Crx^E168d2/+^*^,*Tet-On-hCRX*^ retinae at 3MO (Supplemental Figure 6B), implying a molecular correlation between *CRX* augmentation and robust cone rescue. Notably, endogenous WT CRX expression diminishes with age in *Crx^E168d2/+^* retinae, while induced CRX from the *Tet-On-hCRX* system replenishes overall functional CRX expression in treated retinae (Figure 3A). As a result, the ratio of WT to mutant CRX expression improves in *Crx^E168d2/+^*^,*Tet-On-hCRX*^ retinae at tested ages (Figure 3B). qPCR analysis reveals reduced expression of rod- and cone-enriched genes in *Crx^E168d2/+^* retinae as compared to *WT* controls at various ages (Figure 3C, 3D), with the progressive downregulation of cone-enriched gene expression preceding that of rod-enriched genes. The *Tet-On-hCRX* system boosts expression of these CRX-target genes during the ages of active photoreceptor degeneration, providing the molecular evidence for persistent photoreceptor rescue in *Crx^E168d2/+^*^,*Tet-On-hCRX*^ retinae.

**Figure 3.**
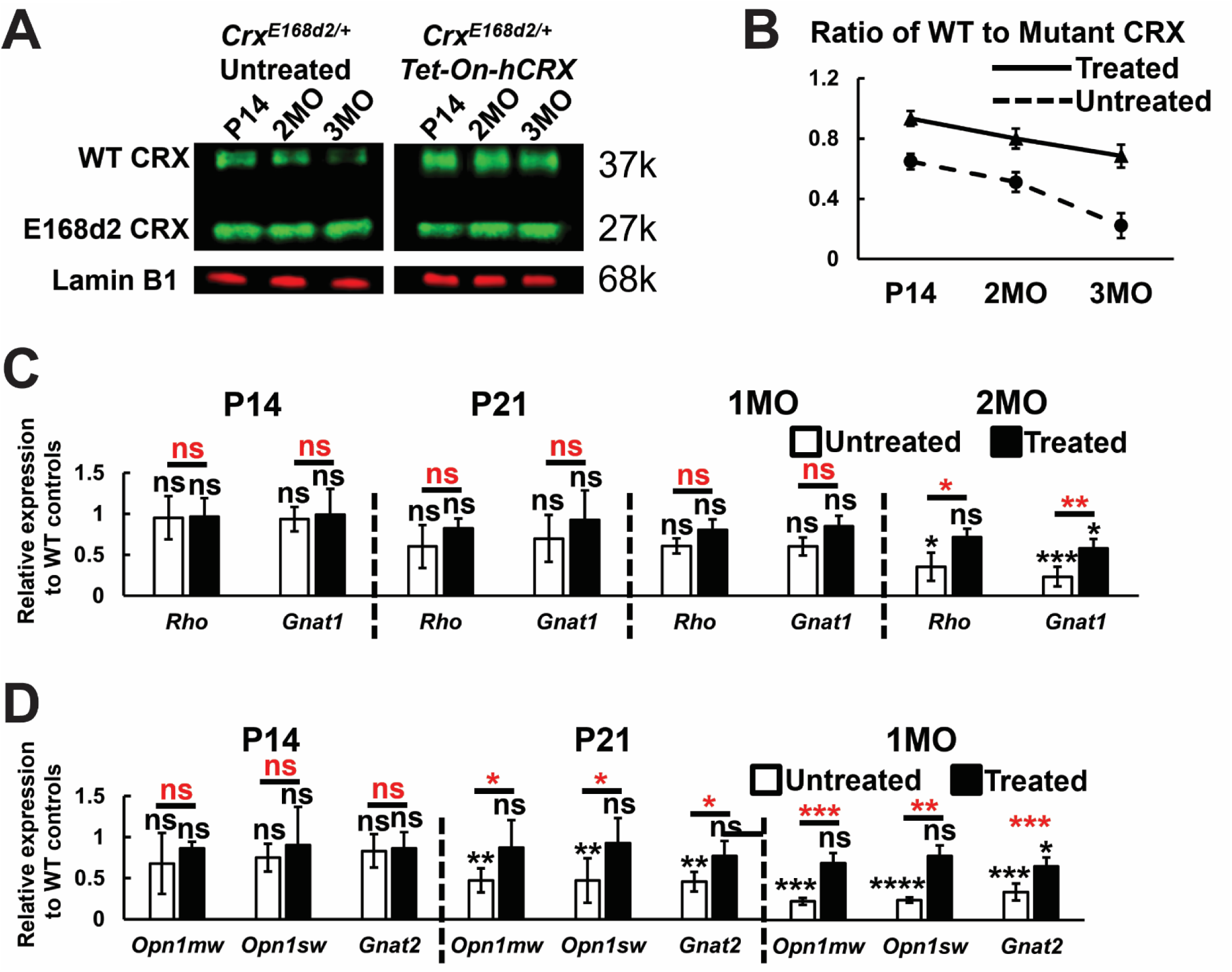
Molecular signature of *CRX* augmentation rescuing *Crx^E168d2/+;Tet-On-hCRX^* retinae. (A) Detection of CRX and Lamin B1 proteins by Western blotting in nuclear extracts of *Crx^E168d2/+^* and *Crx^E168d2/+;Tet-On-hCRX^* retinae at various ages. (B) Ratios of p.WT to p.E168d2 CRX protein at various ages. Ratio is calculated based on fluorescence intensity of Western blotting bands. Error bars represent SD on mean values (n=3). (C) qPCR analysis of rod-enriched genes in *Crx^E168d2/+^* and *Crx^E168d2/+;Tet-On-hCRX^* retinae at various ages. Transcript expression is normalized to *WT* controls. Error bars represent SD on mean values (n=6). (D) qPCR analysis of cone-enriched genes in *Crx^E168d2/+^* and *Crx^E168d2/+;Tet-On-hCRX^* retinae at various ages. Transcript expression is normalized to *WT* controls. Error bars represent SD on mean values (n=6). Statistical analysis by ANOVA happens between *WT*, *Crx^E168d2/+^* and *Crx^E168d2/+;Tet-On-hCRX^* samples for each measurement point. Statistical significance indicated by black asterisks compares with *WT* samples. Statistical significance indicated by red asterisks compares between *Crx^E168d2/+^* and *Crx^E168d2/+;Tet-On-hCRX^* samples. Asterisks (*, **, ***, ****) denote p≤0.05, p≤0.01, p≤0.001 and p≤0.0001 respectively, and ns means not significant.

### *Tet-On-hCRX* rescuing *Crx^E80A/+^* photoreceptors

*CRX^E80A^* mutation belongs to a different pathogenic class of *CRX* mutations, apart from *CRX^E168d2^* mutation. Cone defects appear at a younger age and progress more rapidly in *Crx^E80A/+^* retinae as compared to *Crx^E168d2/+^* retinae (Supplemental figure 1, 2). Hence, a key question asks if the *Tet-On-hCRX* system is effective against the severe yet early-onset disease phenotypes in *Crx^E80A/+^* retinae. Developing RXRγ+ cones are detectable in P0 untreated and *Tet-On-hCRX*-treated *Crx^E80A/+^* retinae (denoted as *Crx^E80A/+^*^,*Tet-On-hCRX*^ in following sections, Supplemental figure 11A), showing comparable cell numbers to those in *WT* retinae (Supplemental figure 11B). Similar to its effect in P0 *Crx^E168d2/+^*^,*Tet-On-hCRX*^ retinae, the *Tet-On-hCRX* system does not increase cone population in neonatal *Crx^E80A/+^*^,*Tet-On-hCRX*^ retinae. Rod development is less affected by *Crx^E80A^* mutation^13^, as seen by similar anti-RHO IHC in untreated and *Crx^E80A/+^*^,*Tet-On-hCRX*^ retinae (Figure 4A). At 1MO, cones are readily detectable in *Crx^E80A/+^*^,*Tet-On-hCRX*^ retinae, in contrast to complete cone loss in untreated retinae (Figure 4B). Notably, a subset of survived cones are found in basal ONL of *Crx^E80A/+^*^,*Tet-On-hCRX*^ retinae (arrows in Figure 4B), whereas cones are usually localized to apical ONL (within 20µm from outer limiting membrane) in *WT* retinae^28–30^. *CRX* augmentation results in moderate improvement in rod-driven ERG responses at high light intensities in 1MO *Crx^E80A/+^*^,*Tet-On-hCRX*^ retinae (Figure 4C, 4D) as compared to untreated retinae. As compared to minimal cone function in untreated retinae, *Crx^E80A/+^*^,*Tet-On-hCRX*^ retinae show significantly boosted cone-driven ERG response (Figure 4E). In summary, these data demonstrate that the *CRX* augmentation via *Tet-On-hCRX* can drive the differentiation and functional maturation of diseased cones in *Crx^E80A/+^* retinae.

**Figure 4.**
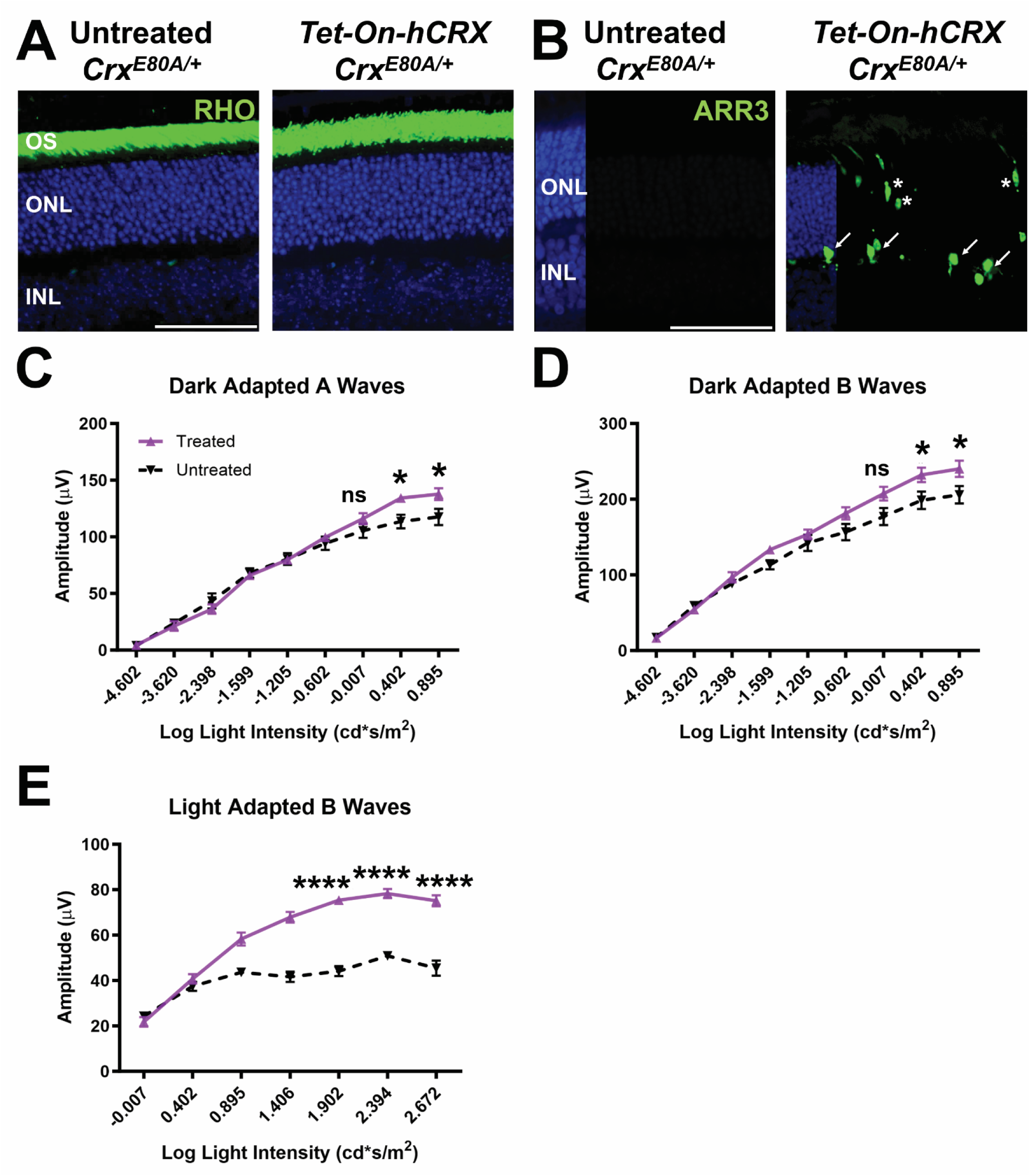
*Tet-On-hCRX*-mediated *CRX* augmentation rescuing *Crx^E80A/+^* retinae. (A) Anti-RHO IHC staining (in green) in *Crx^E80A/+^* and *Crx^E80A/+;Tet-On-hCRX^* retinae at 1MO. Nuclei are stained by DAPI (in blue). Scale bar = 50µm. (B) Anti-ARR3 IHC staining (in green) in *Crx^E80A/+^* and *Crx^E80A/+;Tet-On-hCRX^* retinae at 1MO. Asterisks indicate ARR3+ cones in apical ONL of *Crx^E168d2/+;Tet-On-hCRX^* retinae. Arrows indicate ARR3+ cones in basal ONL. (C) ERG measurements of dark-adapted A waves of *Crx^E80A/+^* and *Crx^E80A/+;Tet-On-hCRX^* retinae mice at 1MO. Error bars represent SD on mean values (n=6). (D) ERG measurements of dark-adapted B waves. (E) ERG measurements of light-adapted B waves. Statistical analysis by pairwise t-test happens between *Crx^E80A/+^* and *Crx^E80A/+;Tet-On-hCRX^* samples for each measurement point. Statistical significance in this figure is indicated by asterisks (*, ****) denoting p≤0.05 and p≤0.0001 respectively, and ns meaning not significant.

Late treatment (initiated at P14) sufficiently rescues cones (Supplemental figure 12A), but at only 50.3% of the efficacy of early treatment (Supplemental figure 12B). Furthermore, late treatment results in only 39.9% of cones localized to apical ONL as compared to early treatment (Supplemental figure 12B). Hence, early treatment facilitates optimal cone migration than late treatment in treated retinae.

### Testing *AAV-*mediated *CRX* augmentation in *Crx* mutant models of cone-rod dystrophies

The excellent efficacy of the *Tet-On-hCRX* system supports the advancement of *AAV*-mediated *CRX* augmentation into preclinical development for *CRX*-associated cone-rod dystrophies. Two choices of promoters, namely *pGrk1* and *pGnb3\**, are selected to drive human *CRX* expression in transduced photoreceptors (Figure 5A). *Grk1* promoter, *pGrk1*, is widely employed to drive *AAV*-mediated gene augmentation in mouse photoreceptors^31,32^. *pGnb3\** has *K50* motifs 1 and 3 replaced by *Q50* motif in *Gnb3* promoter^33^, driving selective gene expression predominantly in photoreceptors. In order to minimize potential toxicity from overexpression, *CRX* augmentation is regulated by the *Tet-Off* system that comprise *tTA* and *TRE* components (Figure 5A). As a safety feature, the *Tet-Off-hCRX* system can be silenced by doxycycline administration if necessary. *AAV2/5*^34,35^ is known to vehicle DNA constructs to photoreceptors. Via *AAV2/5*-meidated delivery, both *pGrk1* and *pGnb3\** promoters efficiently drive a fluorescent reporter in *WT* mouse retinae, showing no signs of overexpression toxicity (Supplemental figure 13A).

**Figure 5.**
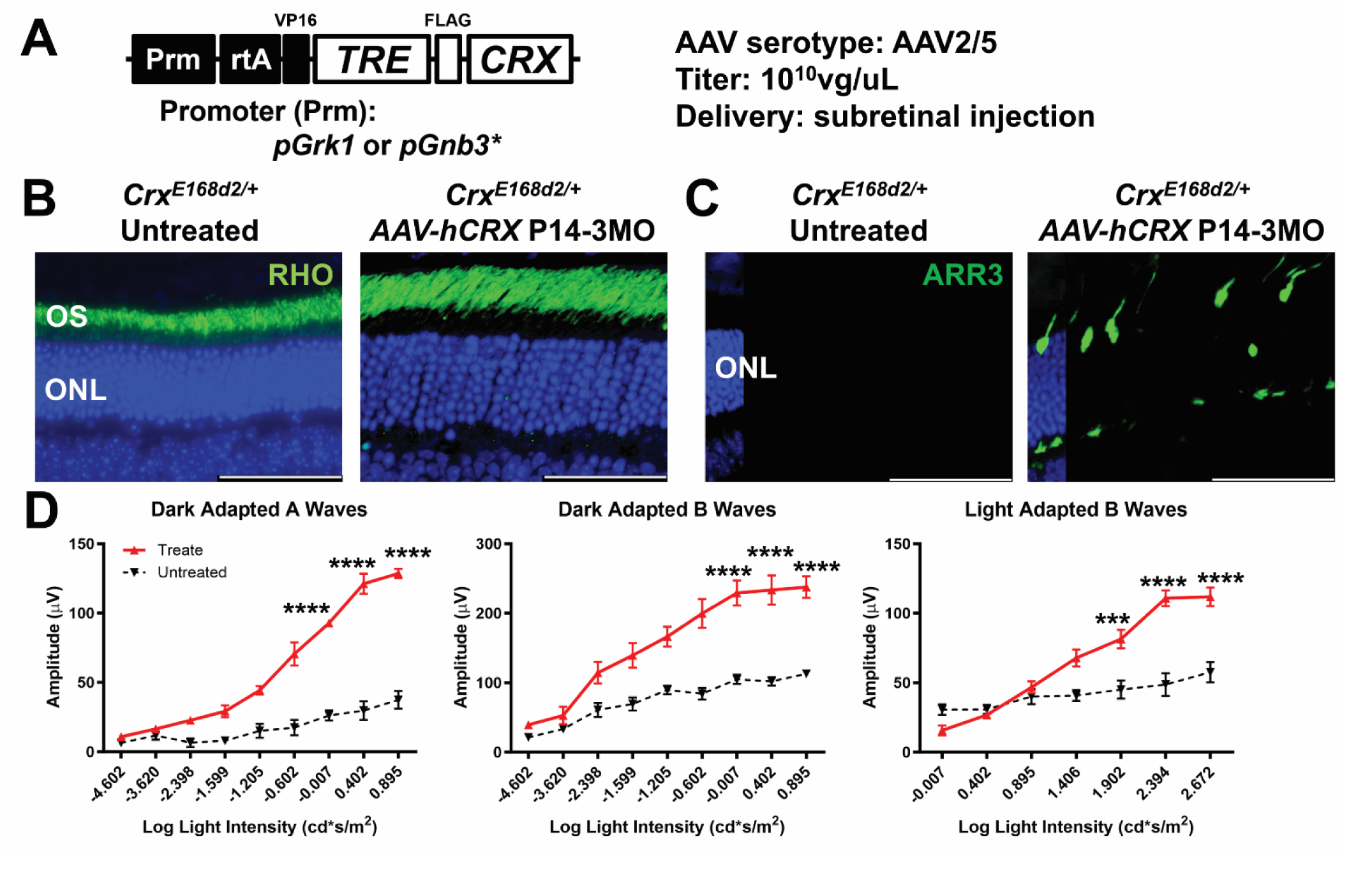
*AAV-hCRX*-mediated *CRX* augmentation rescuing *Crx^E168d2/+^* retinae. (A) Information on AAV construct, choices of promoters, AAV serotype, AAV titer, and delivery. (B) Anti-RHO IHC staining (in green) in *Crx^E168d2/+^* and *Crx^E168d2/+;AAV-hCRX^* retinae at 3MO. Nuclei are stained by DAPI (in blue). Scale bar = 50µm. (C) Anti-ARR3 IHC staining (in green) in *Crx^E168d2/+^* and *Crx^E168d2/+;AAV-hCRX^* retinae at 3MO. (D) ERG measurements of dark-adapted A, B waves and light-adapted B waves of *Crx^E168d2/+^* and *Crx^E168d2/+;AAV-hCRX^* retinae at 3MO. Error bars represent SD on mean values (n=6). Statistical analysis by ANOVA. Statistical significance in this figure is indicated by asterisks (***, ****) denoting p≤0.001 and p≤0.0001 respectively.

*AAV*-mediated *CRX* augmentation is initiated in *Crx^E168d2/+^* retinae at P14, an age point prior to significant cone degeneration (Supplemental figure 1C). As a result, 10^10^vg of *AAV*-*pGnb3*-Tet-Off-hCRX* successfully reinforces RHO localization in OS (Figure 5B) and prolongs cone survival (Figure 5C) in transduced regions at 3MO. *AAV*-*pGrk1-Tet-Off-hCRX* rescues cones with efficacy equal to that of *AAV*-*pGnb3*-Tet-Off-hCRX* (Supplemental figure 13B, 13C). Based on the excellent photoreceptor transduction at neonatal period, *AAV*-*pGnb3*-Tet-Off-hCRX* is chosen to deliver *CRX* augmentation in the following experiments (denoted as *AAV-hCRX*). Two titers, 5X10^8^ and 10^10^vg, are effective in rescuing cones in 2MO *Crx^E168d2/+^* retinae (Supplemental figure 14A), whereas 10^10^vg produces superior therapeutic outcomes (Supplemental figure 14B). Transduction efficiency typically covers more than 30% of photoreceptors. Notably, survived cones are predominantly localized within the transduced regions (Supplemental figure 15). Photoreceptors within the transduced regions display functional recovery, as shown by improved electroretinogram (ERG) responses (Figure 5D).

The important next step is to determine whether *AAV*-mediated *CRX* augmentation follows the same “earlier-the-better” principle observed with the *Tet-On-hCRX* system. *AAV-hCRX* is delivered into *Crx^E168d2/+^* retinae either at P14 (early treatment) or at 1MO (late treatment) (Supplemental figure 16A). Consistent with therapeutic outcomes by the *Tet-Off-hCRX* system, early treatment by *AAV-hCRX* promotes cone survival better than late treatment by 3MO (Supplemental figure 16B).

*AAV-hCRX* achieves effective cone rescue across all tested ages in young adulthood (Figure 6A, 6B, 6C), showing comparable outcomes as the *Tet-On-hCRX* system (Figure 6D). Therefore, the *Tet-On-hCRX* system effectively serves as a robust and reliable proof-of-principle model for testing *AAV-hCRX* in preclinical models of *CRX*-associated cone-rod dystrophies. *CRX* augmentation has been validated by two independent systems as a viable gene therapy strategy for cone rescue in *Crx^E168d2/+^* retinae.

**Figure 6.**
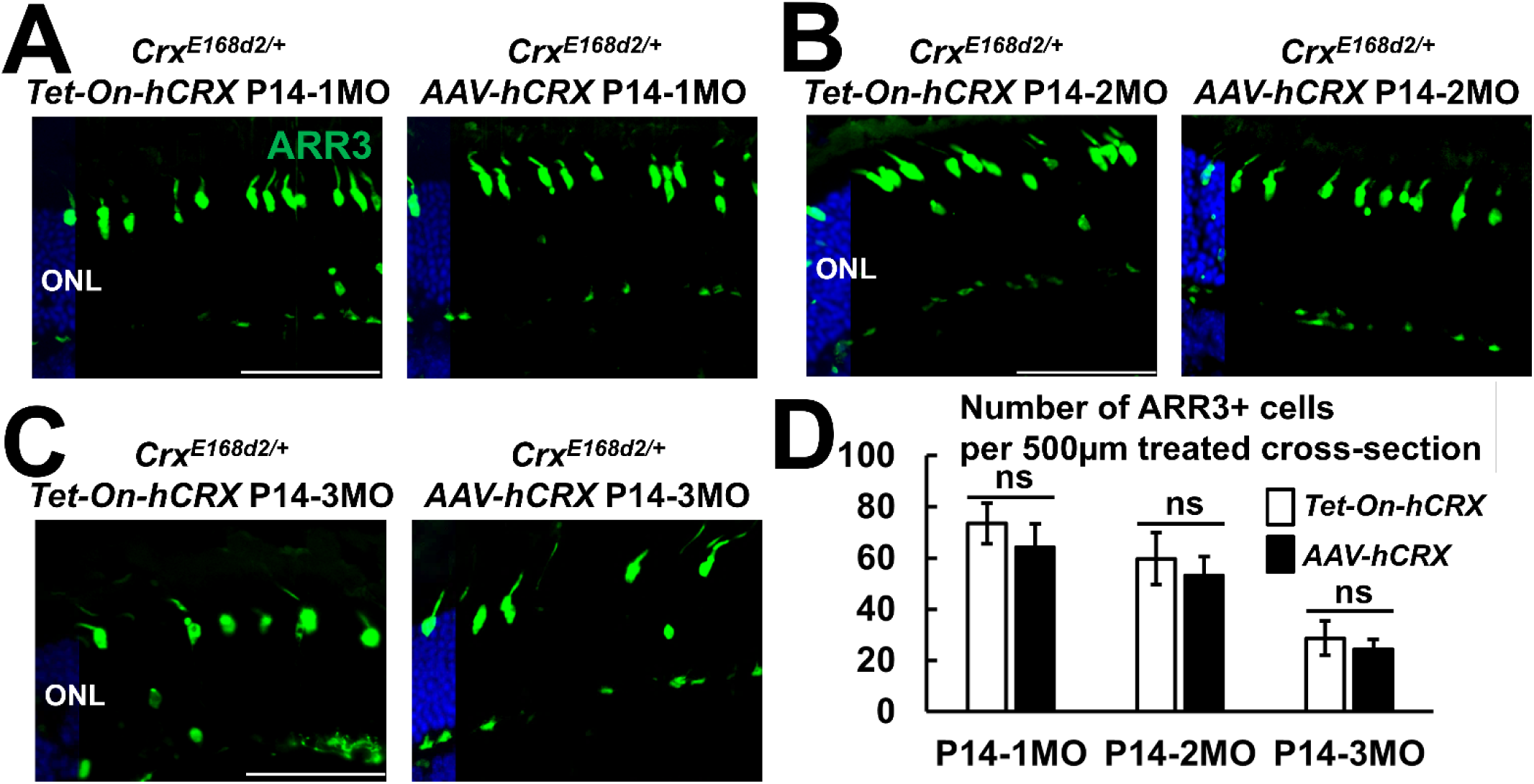
*Tet-On-hCRX* vs *AAV-hCRX* in *Crx^E168d2/+^* retinae. (A-C) Anti-ARR3 IHC staining (in green) in *Crx^E168d2/+^* retinae treated by *Tet-On-hCRX* and *AAV-hCRX* at 1MO, 2MO and 3MO. Nuclei are stained by DAPI (in blue). Scale bars = 50µm. (D) Cone numbers in treated retinae at various ages. Cell count is based on anti-ARR3 IHC staining within treated regions. Error bars represent SD on mean values (n=4). Statistical analysis by pairwise t-test for each age point. Statistical significance in this figure is indicated by ns meaning not significant.

*CRX* augmentation can also foster rod survival after cone loss in *Crx^E168d2/+^* retinae. At 5MO, anti-RHO IHC reveals minimal OS in untreated retinae (Supplemental figure 17A). Interventions initiated at 3MO by either *Tet-Off-hCRX* or *AAV-hCRX* retains OS in treated retinae (Supplemental figure 17B, 17C), demonstrating that *CRX* augmentation can effectively treat late-stage rod degeneration.

*AAV*-mediated *CRX* augmentation also produces promising therapeutic outcomes in *Crx^E80A/+^* retinae. *AAV-hCRX* is applied at P3 (early treatment) or P14 (late treatment). Both regimens promote cone rescue (Supplemental figure 18A), while early treatment yields significantly better therapeutic outcomes (Supplemental figure 18B).

## DISCUSSION

Gene therapy currently represents the leading treatments strategy for inherited retinal diseases. Unlike loss-of-function mutations in structural or enzymatic genes, mutations in transcription factor genes such as *CRX* yield specific therapeutic challenges of precisely regulating target gene expression at critical treatment windows and achieving sufficient functional efficacy. Furthermore, because *CRX^E168d2^* and *CRX^E80A^* are autosomal dominant mutations, the feasible gene therapy must be designed to counteract the detrimental activities of the mutant proteins. Pathogenic analysis has revealed that the insufficient expression of functional CRX disrupts target gene expression in both models, suggesting *CRX* augmentation as a potential therapeutic strategy. In *Crx^E168d2/+^* retinae, *CRX* augmentation prolongs photoreceptor survival, most notably for cones, in young adulthood. *CRX* augmentation upregulates the expression of phototransduction genes in treated retinae, thus boosting functional recovery. The rescue effect is sustained into later adulthood. In *Crx^E80A/+^* retinae, *CRX* augmentation facilitates cone differentiation and supports functional maturation. Considering the early-onset cone loss, *CRX* augmentation promotes structural development, phototransduction assembly, and proper cell migration in a subset of treated cones. All in all, this study successfully validates *CRX* augmentation as a viable therapeutic strategy for *CRX*-associated IRDs.

The success of *CRX* augmentation relies on intervention timing, following the ‘earlier-the-better’ principle for therapeutic efficacy. Three important conclusions can be made. Firstly, early intervention targets the degenerative mechanisms at the initial onset. Untreated *Crx^E168d2/+^* retinae show misregulated photoreceptor gene expression, especially for phototransduction genes, leading to progressive and persistent functional deterioration. In treated *Crx^E168d2/+^* retinae, CRX augmentation upregulates photoreceptor gene expression and enhances phototransduction pathways, which in turn promotes photoreceptor survival. Untreated *Crx^E80A/+^* retinae show disrupted cone differentiation due to insufficient expression of p.WT CRX for binding activity at target genes. *CRX* augmentation drives the timely cone differentiation by restoring the complex regulatory gene network. Secondly, functional recovery depends on the intervention timing. When *CRX* augmentation operates at optimal capacity (10^10^vg/uL), early treatment produces significantly better improvements in visual functions than late treatment. Lastly, late treatment can still achieve positive therapeutic outcomes, suggesting a degree of plasticity in the treatment window.

The *Tet-On-hCRX* system serves as a valuable preclinical platform for evaluating the therapeutic effectiveness of *CRX* augmentation. Firstly, *Tet-On-hCRX* can activate robust *CRX* augmentation at early neonatal ages. This feature enables the faithful evaluation of *CRX* augmentation in models of early-onset IRDs such as LCA or cone-rod dystrophy. In contrast, *AAV*-mediated gene therapy typically requires up to two weeks to reach peak transgene expression^36,37^. Secondly, *Tet-On-hCRX* induces pan-photoreceptor *CRX* augmentation, which permits the comprehensive and unbiased assessments of functional recovery in treated retinae. Thirdly, *Tet-On-hCRX* induces consistent expression of *CRX* augmentation in treated photoreceptors, which ensures accurate molecular analysis. Lastly, *Tet-On-hCRX* induces steady *CRX* augmentation among biological replicates, which minimizes experimental variation and invasive operations. On the other hands, *AAV*-mediated gene therapy often shows variable transgene expression due to inconsistency in viral copy number and transduction efficiency^38,39^.

*AAV*-mediated CRX augmentation is a promising therapeutic approach for *CRX*-associated cone-rod dystrophies. Firstly*, AAV2/5* can readily transduce diseased cones in mouse models even at neonatal period. Secondly, in addition to the commonly used *pGrk1*, this study has validated *pGnb3\** as a functional promoter for driving transgene expression in developing and mature photoreceptors. In addition, *pGnb3\** may be minimally affected by CRX mutant proteins. Lastly, none of the administered titers causes detectable overexpression toxicity in treated retinae, confirming the biosafety of *AAV*-mediated *CRX* augmentation regimen.

*CRX* augmentation does not halt the degeneration but significantly alters the degenerative trajectory. The primary goal of this study has been successfully achieved that *CRX* augmentation prolongs cone survival in *Crx^E168d2/+^* and *Crx^E80A/+^* retinae. *CRX* augmentation targets the pathogenic mechanisms rather than merely rectifying the cell death program in diseased photoreceptors. Moreover, mutant proteins are still present in the treated retinae, which ultimately drives the degenerative process. The degenerative mechanisms in treated retinae remain unclear. In terms of designing more effective therapeutic strategies, escalating beyond the currently applied titer (10^10^vg/uL) risks causing overexpression toxicity and severe immune responses. Future studies will develop complementary genetic approaches such as allele-specific CRISPR/Cas9 knockout or shRNA-mediated knockdown to reduce mutant *CRX* expression.

Beyond the therapeutic implications, this study also addresses critical questions in photoreceptor biology. Firstly, *CRX* overexpression does not alter cone birth or population, suggesting that CRX only regulates photoreceptor fate specification rather than the overall photoreceptor composition. Secondly, *Crx^-/-^*retinae lack OS structure or photoreceptor function^3^, whereas untreated *Crx^E168d2/+^* and *Crx^80A/+^* retinae, having ≤50% of *WT* Crx expression, retain normal retinal morphology in young adulthood but have minimal photoreceptor function. *CRX* augmentation promotes functional recovery in treated retinae of both models. On the other hand, *CRX* overexpression in *WT* retinae does not promote photoreceptor function. These findings altogether suggest that misregulated gene expression causes functional loss prior to massive photoreceptor degeneration; Sufficient *CRX* expression is indispensable for maintaining photoreceptor function but does not drive the phototransduction cascade by itself. Future studies will define the quantitative correlation between *CRX* expression and the efficiency of the phototransduction cascade. Lastly, cone differentiation depends more on *WT CRX* expression and its protein binding activity than rod differentiation^23^. *CRX* augmentation facilitates the migration of some cones to the apical ONL, implying the role of *CRX* expression in cone migration during early development. Future studies include multi-omics analysis to elucidate alterations in CRX binding and chromatin remodeling by *CRX* augmentation, revealing in-depth therapeutic mechanisms.

## ETHICS STATEMENT

The animal study was reviewed and approved by Washington University in St. Louis Institutional Animal Care and Use Committee.

## AUTHOR CONTRIBUTIONS

CS and SC designed experiments. CS performed the procedures of *AAV* injection, IHC staining, imaging, ERG, Western blotting, RNA sample preparation, and qPCR. MF performed OKR and PLR. CS and SC prepared the manuscript. All authors edited and revised the manuscript.

## ACKNOWLEDGMENTS

Authors are grateful to Ms. Mingyan Yang and Dr. Guangyi Ling for technical assistance. The following grants have supported this study: Knights Templar Pediatric Ophthalmology Career-Starter Research Grant (CS), VitreoRetinal Surgery Foundation Research Fellowship (CS), McDonnell Center for Cellular and Molecular Neurobiology Fellowship (CS), R01 EY012543 (SC), R01 EY032136 (SC), Widłak Family CRX Research Fund (SC), Kompania Gornicza CRX Research Fund (SC), R01 EY027411 (DK), R01 EY034001 (DK), R01 EY026978 (DK), P30 EY002687 (WUSTL DOVS), Research to Prevent Blindness (WUSTL DOVS).

