## Supplemental Figures and Tables for "Preclinical *CRX* augmentation therapies for *CRX*-associated autosomal dominant cone-rod dystrophies": CRX Augmentation MS Supplemental Figure.pdf

Supplemental figure 1

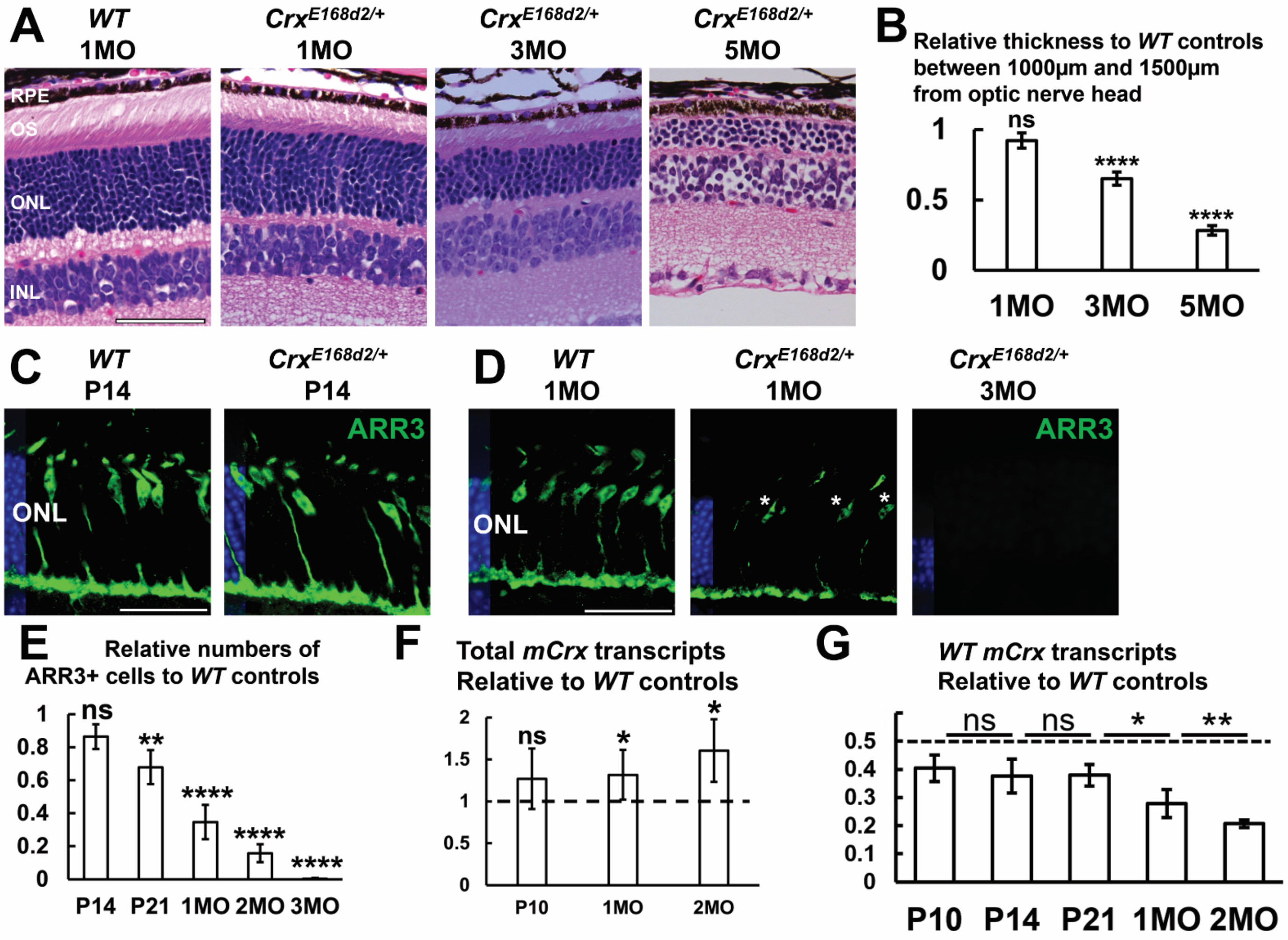

Supplemental figure 1. Photoreceptor degeneration in *Crx*$^{E168d2/+}$ retinae. (A) H&E cross-section staining of *WT* retinae at 1MO and *Crx*$^{E168d2/+}$ retinae at 1MO, 3MO and 5MO. RPE: retinal pigment epithelium. OS: outer segment. ONL: outer nuclear layer. INL: inner nuclear layer. Scale bar = 50µm. (B) Relative ONL thickness to *WT* controls at 1MO, 3MO and 5MO. ONL thickness is measured between 1000 and 1500µm from optical nerve head. Error bars represent SD on mean values (n=4). Statistical analysis by t-test happens between *Crx*$^{E168d2/+}$ and *WT* samples at each age point. (C) Anti-ARR3 IHC staining (in green) in *Crx*$^{E168d2/+}$ and *WT* retinae at P14. Nuclei are stained by DAPI (in blue). Scale bar = 50µm. (D) Anti-ARR3 IHC staining (in green) in *WT* retinae at 1MO and *Crx*$^{E168d2/+}$ retinae at 1MO and 3MO. Asterisks indicate examples of ARR3-labelled cells in 1MO *Crx*$^{E168d2/+}$ retinae. (E) Relative cone numbers to *WT* controls at P14, P21, 1MO, 2MO and 3MO. Cell count is based on anti-ARR3 IHC staining between 500 to 1500µm from optical nerve head. Error bars represent SD on mean values (n=4). Statistical analysis by t-test happens between *Crx*$^{E168d2/+}$ and *WT* samples at each age point. (F) qPCR analysis of total mouse *Crx* (*mCrx*) transcript expression in *Crx*$^{E168d2/+}$ and *WT* retinae at P10, 1MO and 2MO. Error bars represent SD on mean values (n=5). Statistical analysis by t-test happens between *Crx*$^{E168d2/+}$ and *WT* samples at each age point. (G) qPCR analysis of *WT mCrx* transcript expression in *Crx*$^{E168d2/+}$ and *WT* retinae at P10, P14, P21, 1MO and 2MO. Error bars represent SD on mean values (n=5). Statistical analysis by ANOVA happens between *Crx*$^{E168d2/+}$ and *WT* samples at each age point as well as mutants of different age points. Statistical significance in this figure is indicated by asterisks (*, **, ****) denoting p≤0.05, p≤0.01, and p≤0.0001 respectively, and ns meaning not significant.

Supplemental figure 2

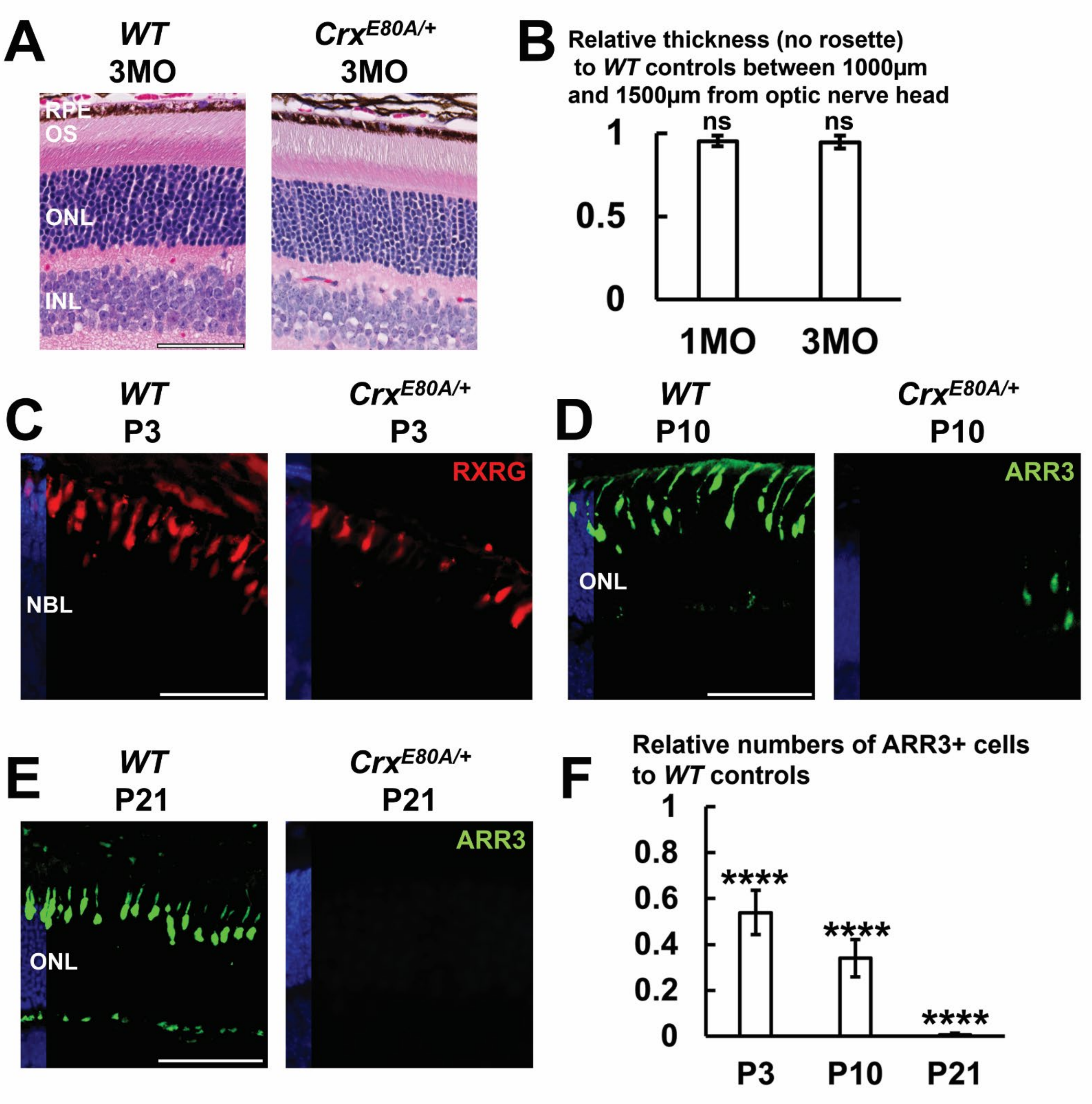

Supplemental figure 2. Photoreceptor degeneration in *Crx*$^{E80A/+}$ retinae. (A) H&E cross-section staining of *WT* and *Crx*$^{E80A/+}$ retinae at 3MO. Scale bar = 50µm. (B) Relative ONL thickness to *WT* controls at 1MO and 3MO. ONL thickness is measured between 1000 and 1500µm from optical nerve head. Error bars represent SD on mean values (n=4). (C) Anti- RXRγ IHC staining (in red) in *WT* and *Crx*$^{E80A/+}$ retinae at P3. Nuclei are stained by DAPI (in blue). Scale bar = 50µm. NBL: neuroblast layer. (D) Anti-ARR3 IHC staining (in green) in *WT* and *Crx*$^{E80A/+}$ retinae at P10. (D) Anti-ARR3 IHC staining (in green) in *WT* and *Crx*$^{E80A/+}$ retinae at P21. (D) Relative cone numbers to *WT* controls at P3, P10 and P21. Cell count is based on anti- RXRγ (P3) and anti-ARR3 (P10 and P21) IHC staining. Error bars represent SD on mean values (n=4). Statistical analysis by t-test happens between *Crx*$^{E80A/+}$ and *WT* samples at each age point. Statistical significance in this figure is indicated by asterisks (****) denoting p≤0.0001 and ns meaning not significant.

Supplemental figure 3

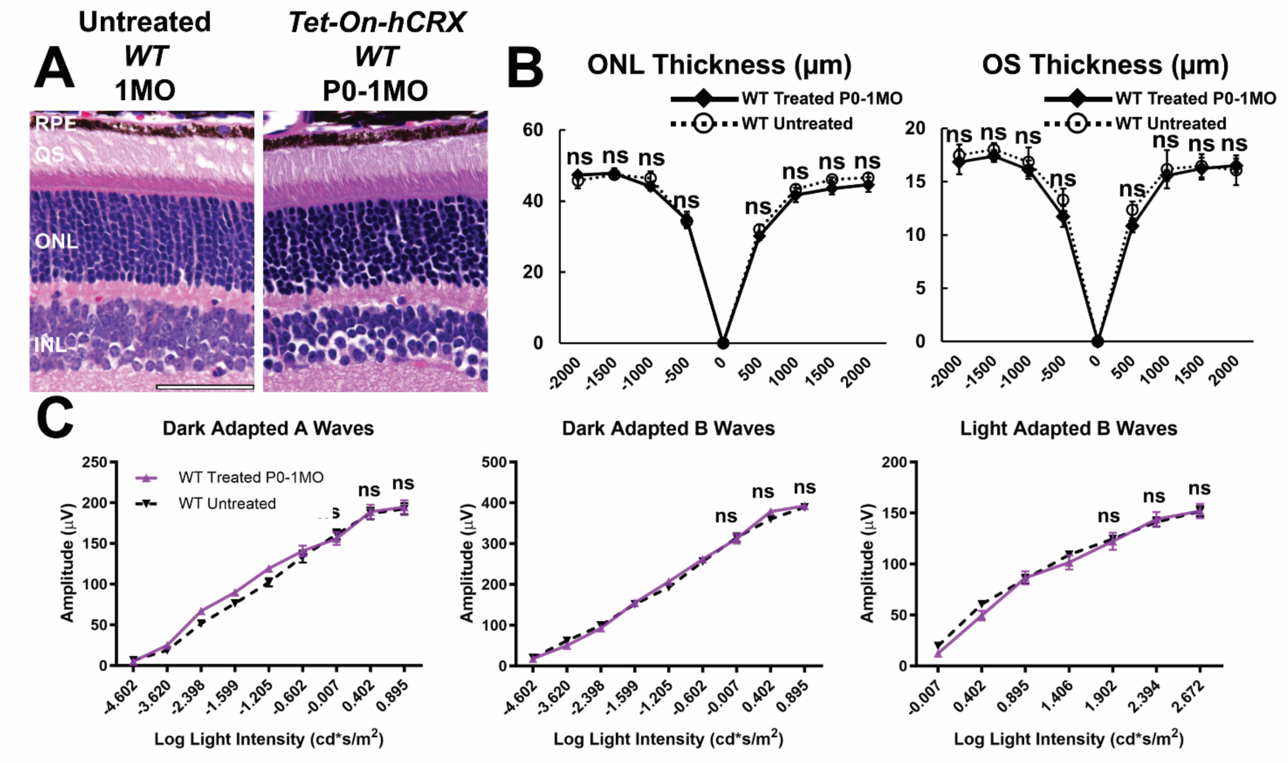

Supplemental figure 3. Safety of *Tet-On-hCRX* system. (A) H&E cross-section staining of untreated *WT* and *WT*$^{Tet\text{-}On\text{-}hCRX}$ retinae at 1MO. Scale bar = 50μm. (B) ONL and OS thickness of untreated *WT* and *WT*$^{Tet\text{-}On\text{-}hCRX}$ retinae at 1MO. Error bars represent SD on mean values (n=4). (C) Electroretinogram (ERG) measurements of dark-adapted A waves, dark-adapted B waves and light-adapted B waves of untreated *WT* and *WT*$^{Tet\text{-}On\text{-}hCRX}$ mice at 1MO. Error bars represent SD on mean values (n=4). Statistical analysis by pairwise t-test happens between untreated *WT* and *WT*$^{Tet\text{-}On\text{-}hCRX}$ samples for each measurement point. Statistical significance in this figure is indicated by ns meaning not significant.

Supplemental figure 4

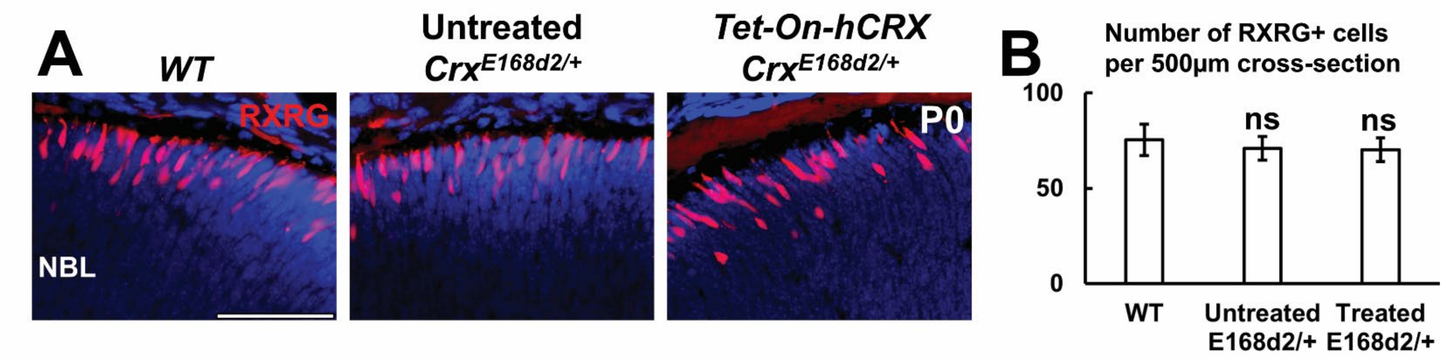

Supplemental figure 4. $Crx^{E168d2/+}$ retinae at P0. (A) Anti- RXRγ IHC staining (in red) in *WT*, $Crx^{E168d2/+}$, $Crx^{E168d2/+;Tet\text{-}On\text{-}hCRX}$ retinae at P0. Nuclei are stained by DAPI (in blue). Scale bar = 50μm. (B) Cone numbers in *WT*, $Crx^{E168d2/+}$, $Crx^{E168d2/+;Tet\text{-}On\text{-}hCRX}$ retinae at P0. Cell count is based on anti-RXRγ IHC staining between 500 to 1000μm from optical nerve head. Error bars represent SD on mean values (n=4). Statistical analysis by ANOVA happens between all groups. Statistical significance in this figure is indicated by ns meaning not significant.

Supplemental figure 5

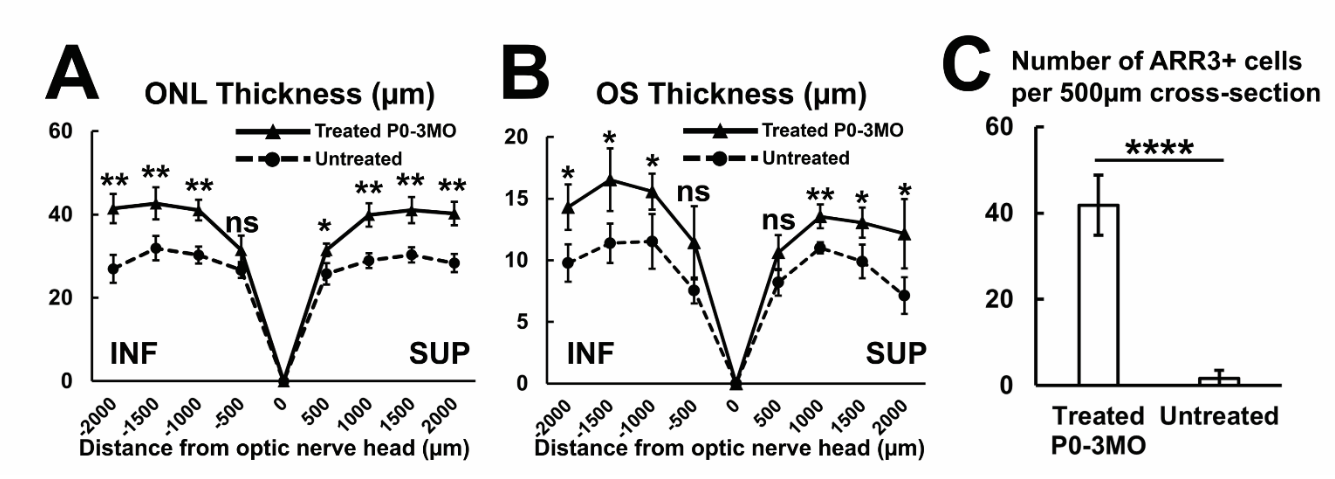


\

Supplemental figure 5. Improved ONL and OS thickness of $Crx^{E168d2/+;Tet\text{-}On\text{-}hCRX}$ retinae at 3MO. (A) ONL thickness of $Crx^{E168d2/+}$ and $Crx^{E168d2/+;Tet\text{-}On\text{-}hCRX}$ retinae at 3MO. Error bars represent SD on mean values (n=4). (B) OS thickness of $Crx^{E168d2/+}$ and $Crx^{E168d2/+;Tet\text{-}On\text{-}hCRX}$ retinae at 3MO. Error bars represent SD on mean values (n=4). (C) Cone numbers in $Crx^{E168d2/+}$ and $Crx^{E168d2/+;Tet\text{-}On\text{-}hCRX}$ retinae at 3MO. Cell count is based on anti-ARR3 IHC staining between 1000 to 1500μm from optical nerve head. Error bars represent SD on mean values (n=4). Statistical analysis by pairwise t-test happens between $Crx^{E168d2/+}$ and $Crx^{E168d2/+;Tet\text{-}On\text{-}hCRX}$ samples for each measurement point. Statistical significance in this figure is indicated by asterisks (*, **, ****) denoting $p \leq 0.05$, $p \leq 0.01$, and $p \leq 0.0001$ respectively, and ns meaning not significant.

Supplemental figure 6

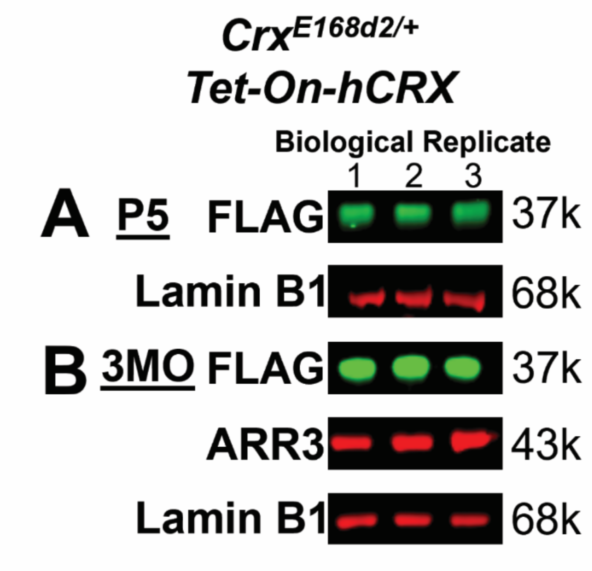

Supplemental figure 6. *CRX* augmentation detected by Western blotting. (A) Western blotting showing FLAG bands with nuclear extracts of P5 $Crx^{E168d2/+;Tet\text{-}On\text{-}hCRX}$ retinae. (B) Western blotting showing FLAG and ARR3 bands with nuclear extracts of 3MO $Crx^{E168d2/+;Tet\text{-}On\text{-}hCRX}$ retinae. 3 biological replicates are shown.

Supplemental figure 7

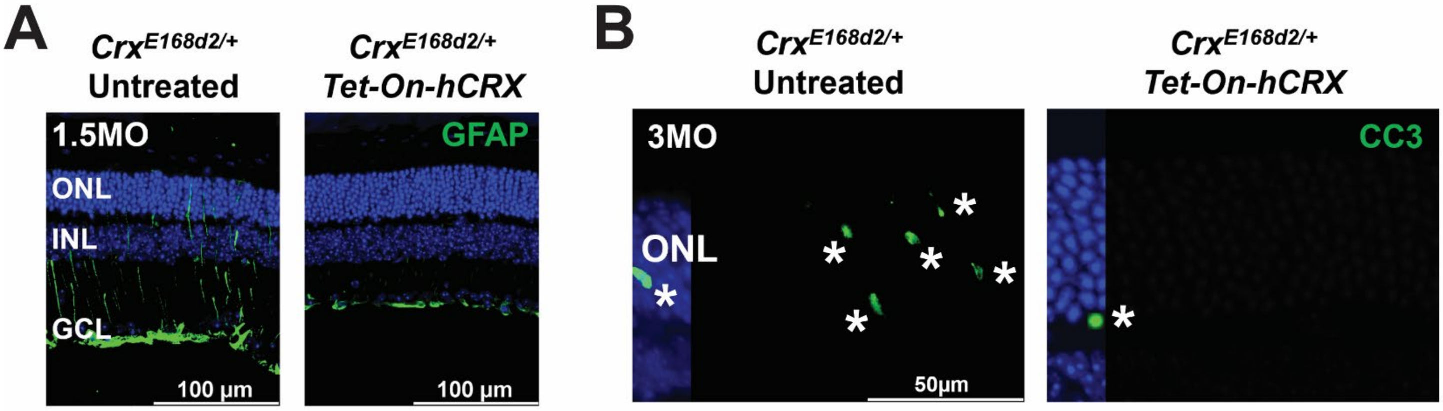

Supplemental figure 7. Glial activation and apoptosis in *Crx*$^{E168d2/+;Tet\text{-}On\text{-}hCRX}$ retinae. (A) Detection of glial fibrillary acidic protein (GFAP) by IHC staining (in green) in *Crx*$^{E168d2/+}$ and *Crx*$^{E168d2/+;Tet\text{-}On\text{-}hCRX}$ retinae at 1.5MO. (B) Detection Cleaved caspase 3 (CC3) IHC staining (in green) in *Crx*$^{E168d2/+}$ and *Crx*$^{E168d2/+;Tet\text{-}On\text{-}hCRX}$ retinae at 3MO. Asterisks indicate examples of CC3-labelled cells. Nuclei are stained by DAPI (in blue). Scale bars represent 100μm in A and 50μm in B.

Supplemental figure 8

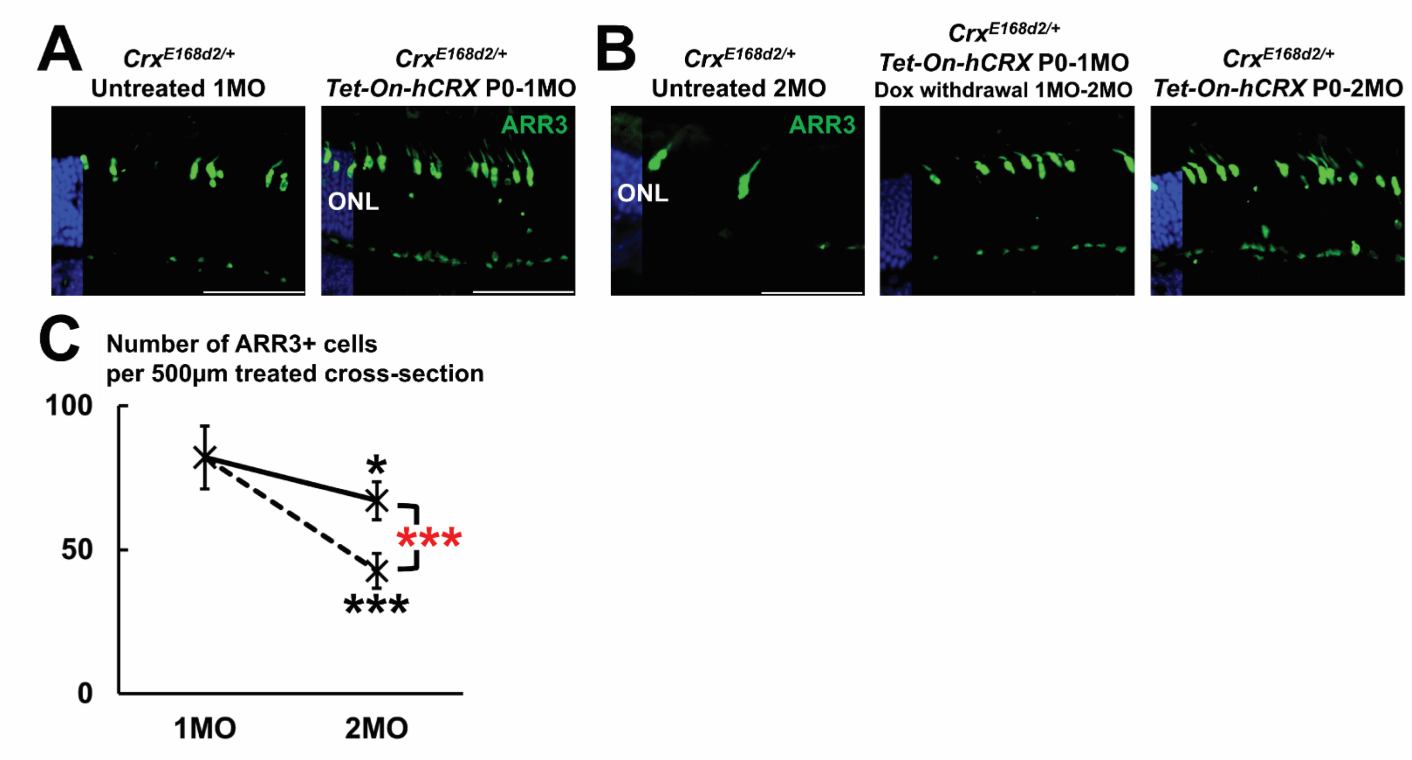

Supplemental figure 8. Doxycycline withdrawal in $Crx^{E168d2/+;Tet\text{-}On\text{-}hCRX}$ retinae. (A) Anti-ARR3 IHC staining (in green) in $Crx^{E168d2/+}$ and $Crx^{E168d2/+;Tet\text{-}On\text{-}hCRX}$ retinae at 1MO. Nuclei are stained by DAPI (in blue). Scale bars = 50µm. (B) Anti-ARR3 IHC staining (in green) in $Crx^{E168d2/+}$, dox-withdrawn $Crx^{E168d2/+;Tet\text{-}On\text{-}hCRX}$ and $Crx^{E168d2/+;Tet\text{-}On\text{-}hCRX}$ (i.e. continuous *Tet-On-hCRX*) retinae at 2MO. Dox withdrawal happens to $Crx^{E168d2/+;Tet\text{-}On\text{-}hCRX}$ at 1MO and the *Tet-On-hCRX* system remains off until sample harvest at 2MO. (C) Cone numbers in tested retinae at 1MO and 2MO. Cell count is based on anti-ARR3 IHC staining between 1000 to 1500µm from optical nerve head. Error bars represent SD on mean values (n=4). Statistical analysis is performed with ANOVA. Statistical significance indicated by black asterisks compares between 2MO samples and 1MO $Crx^{E168d2/+;Tet\text{-}On\text{-}hCRX}$ retinae. Statistical significance indicated by red asterisks compares between 2MO samples. Asterisks (**, ***) denote p≤0.01 and p≤0.001 respectively.

Supplemental figure 9

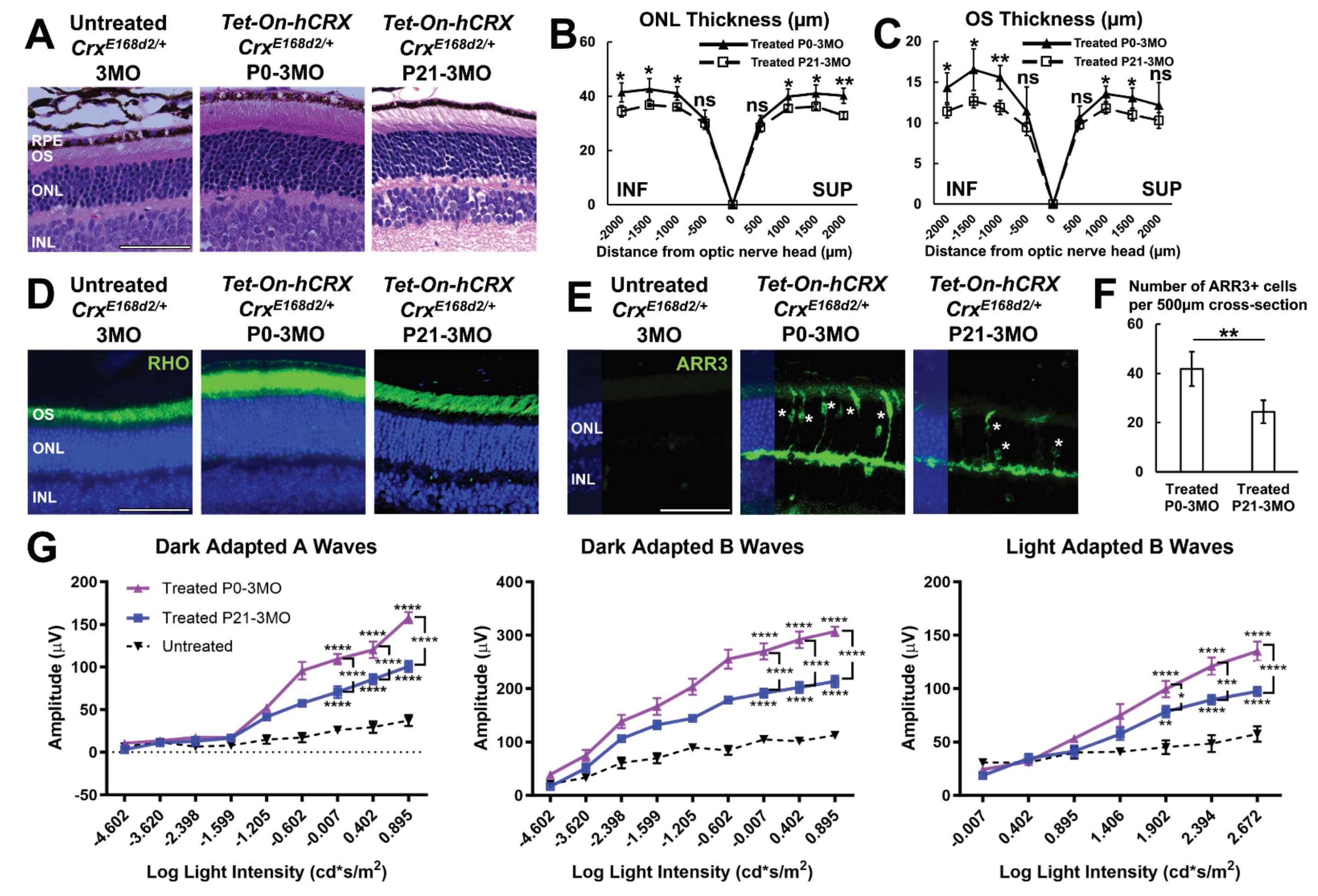

Supplemental figure 9. Early vs late treatment by *Tet-On-hCRX*-mediated *CRX* augmentation in *Crx*$^{E168d2/+}$ retinae. Early treatment means P0-3MO, late treatment means P21-3MO. (A) H&E cross-section staining of samples treated by different regimes at 3MO. Scale bar = 50µm. (B) ONL thickness of samples treated by different regimes at 3MO. Error bars represent SD on mean values (n=4). (C) OS thickness of samples treated by different regimes at 3MO. Error bars represent SD on mean values (n=4). (D) Anti-RHO IHC staining (in green) of samples treated by different regimes at 3MO. Nuclei are stained by DAPI (in blue). (E) Anti-ARR3 IHC staining (in green) of samples treated by different regimes at 3MO. Asterisks indicate ARR3+ cones. (F) Cone numbers in samples treated by different regimes at 3MO. Cell count is based on anti-ARR3 IHC staining between 1000 to 1500µm from optical nerve head. Error bars represent SD on mean values (n=4). (G) ERG measurements of dark-adapted A, B waves and light-adapted B waves of samples treated by different regimes at 3MO. Error bars represent SD on mean values (n=6). Statistical analysis by pairwise t-test for B, C and F and by ANOVA for G. Statistical significance in this figure is indicated by asterisks (*, **, ***, ****) denoting $p \leq 0.05$, $p \leq 0.01$, $p \leq 0.001$ and $p \leq 0.0001$ respectively, and ns meaning not significant.

Supplemental figure 10

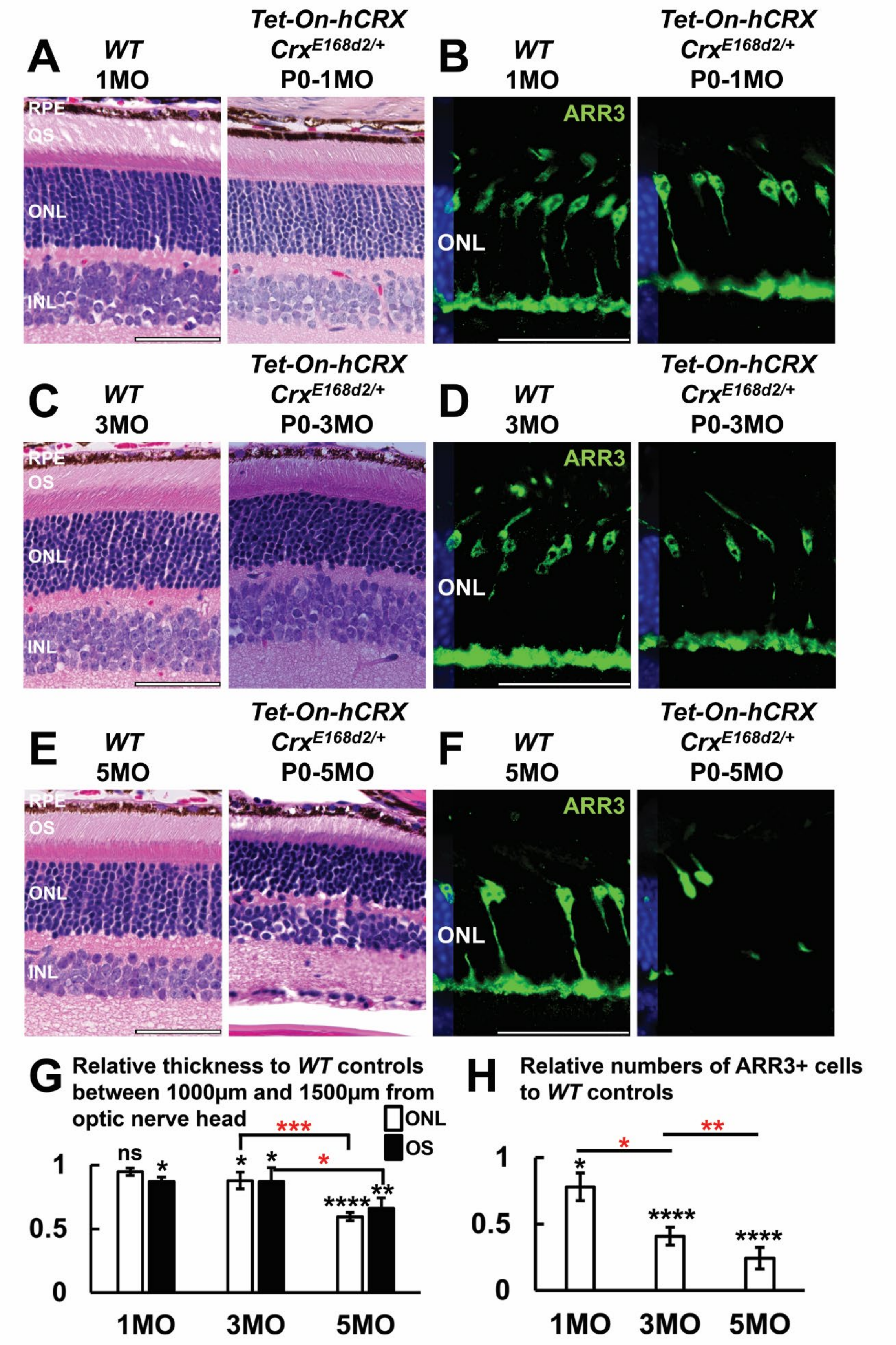

Supplemental figure 10. Comparison between *WT* and $Crx^{E168d2/+;Tet\text{-}On\text{-}hCRX}$ retinae. (A) H&E cross-section staining of *WT* and $Crx^{E168d2/+;Tet\text{-}On\text{-}hCRX}$ retinae at 1MO. Scale bar = 50μm. (B) Anti-ARR3 IHC staining (in green) in *WT* and $Crx^{E168d2/+;Tet\text{-}On\text{-}hCRX}$ retinae at 1MO. Nuclei are stained by DAPI (in blue). Scale bar = 50μm. (C) H&E cross-section staining of *WT* and $Crx^{E168d2/+;Tet\text{-}On\text{-}hCRX}$ retinae at 3MO. (D) Anti-ARR3 IHC staining (in green) in *WT* and $Crx^{E168d2/+;Tet\text{-}On\text{-}hCRX}$ retinae at 3MO. (E) H&E cross-section staining of *WT* and $Crx^{E168d2/+;Tet\text{-}On\text{-}hCRX}$ retinae at 5MO. (F) Anti-ARR3 IHC staining (in green) in *WT* and $Crx^{E168d2/+;Tet\text{-}On\text{-}hCRX}$ retinae at 5MO. (G) ONL and OS thickness of *WT* and $Crx^{E168d2/+;Tet\text{-}On\text{-}hCRX}$ retinae between 1000 to 1500μm at various ages. Error bars represent SD on mean values (n=4). (H) Cone numbers in $Crx^{E168d2/+;Tet\text{-}On\text{-}hCRX}$ retinae relative to *WT* controls at various ages. Cell count is based on anti-ARR3 IHC staining between 1000 to 1500μm from optical nerve head. Error bars represent SD on mean values (n=4). Statistical analysis by ANOVA. Statistical significance indicated by black asterisks compares with *WT* and $Crx^{E168d2/+;Tet\text{-}On\text{-}hCRX}$ retinae. Statistical significance indicated by red asterisks compares $Crx^{E168d2/+;Tet\text{-}On\text{-}hCRX}$ retinae of different ages. *, **, **** denotes $p \leq 0.05$, $p \leq 0.01$, and $p \leq 0.0001$ respectively, and ns means not significant.

Supplemental figure 11

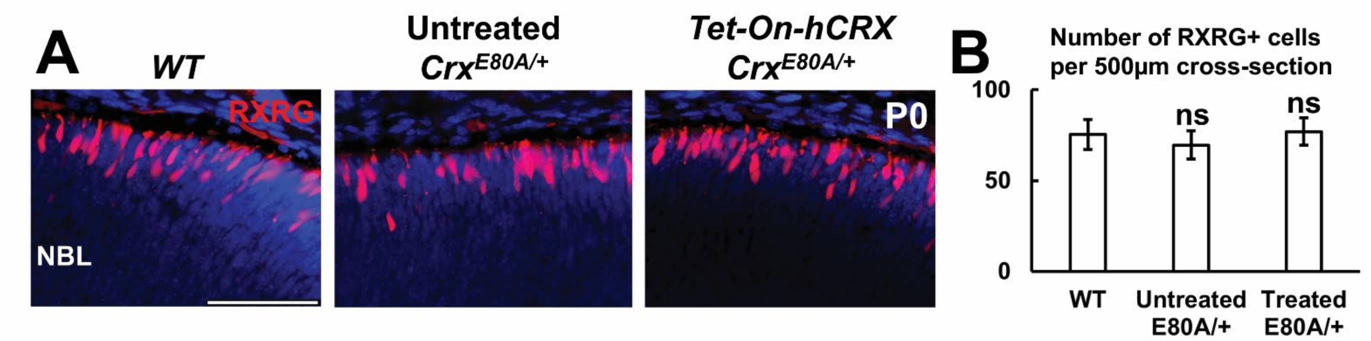

Supplemental figure 11. $Crx^{E80A/+}$ retinae at P0. (A) Anti-RXRG IHC staining (in red) in *WT*, $Crx^{E80A/+}$, $Crx^{E80A/+;Tet\text{-}On\text{-}hCRX}$ retinae at P0. Nuclei are stained by DAPI (in blue). Scale bar = 50μm. (B) Cone numbers in *WT*, $Crx^{E80A/+}$, $Crx^{E80A/+;Tet\text{-}On\text{-}hCRX}$ retinae at P0. Cell count is based on anti-RXRG IHC staining between 500 to 1000μm from optical nerve head. Error bars represent SD on mean values (n=4). Statistical analysis by ANOVA happens between all groups. Statistical significance in this figure is indicated by ns meaning not significant.

Supplemental figure 12

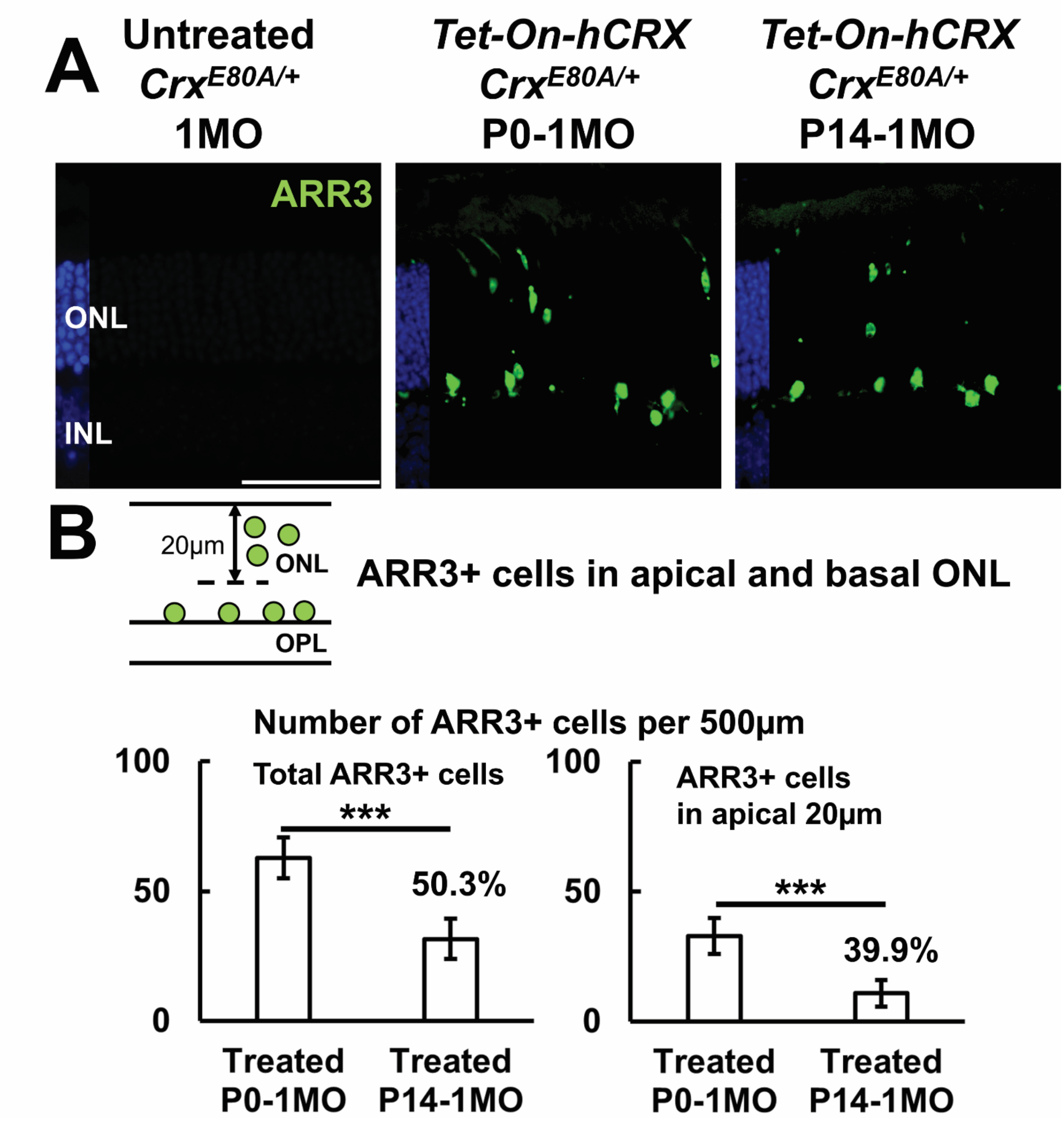

Supplemental figure 12. Early vs late treatment by *Tet-On-hCRX*-mediated *CRX* augmentation in *Crx*$^{E80A/+}$ retinae. Early treatment means P0-1MO, late treatment means P14-1MO. (A) Anti-ARR3 IHC staining (in green) in samples treated by different regimes at 1MO. Nuclei are stained by DAPI (in blue). Scale bar = 50μm. (B) Illustration and cell counts of cones in apical and basal ONL. Cell count is based on anti-ARR3 IHC staining between 1000 to 1500μm, tallying total cell numbers (left plot) and cells in apical ONL (right plot). Percentages describe the ratios of the average cell numbers rescued by late treatment to those by early treatment. Error bars represent SD on mean values (n=4). Statistical analysis by pairwise t-test between samples treated by different regimes. Asterisks (***) denote $p \leq 0.001$.

Supplemental figure 13

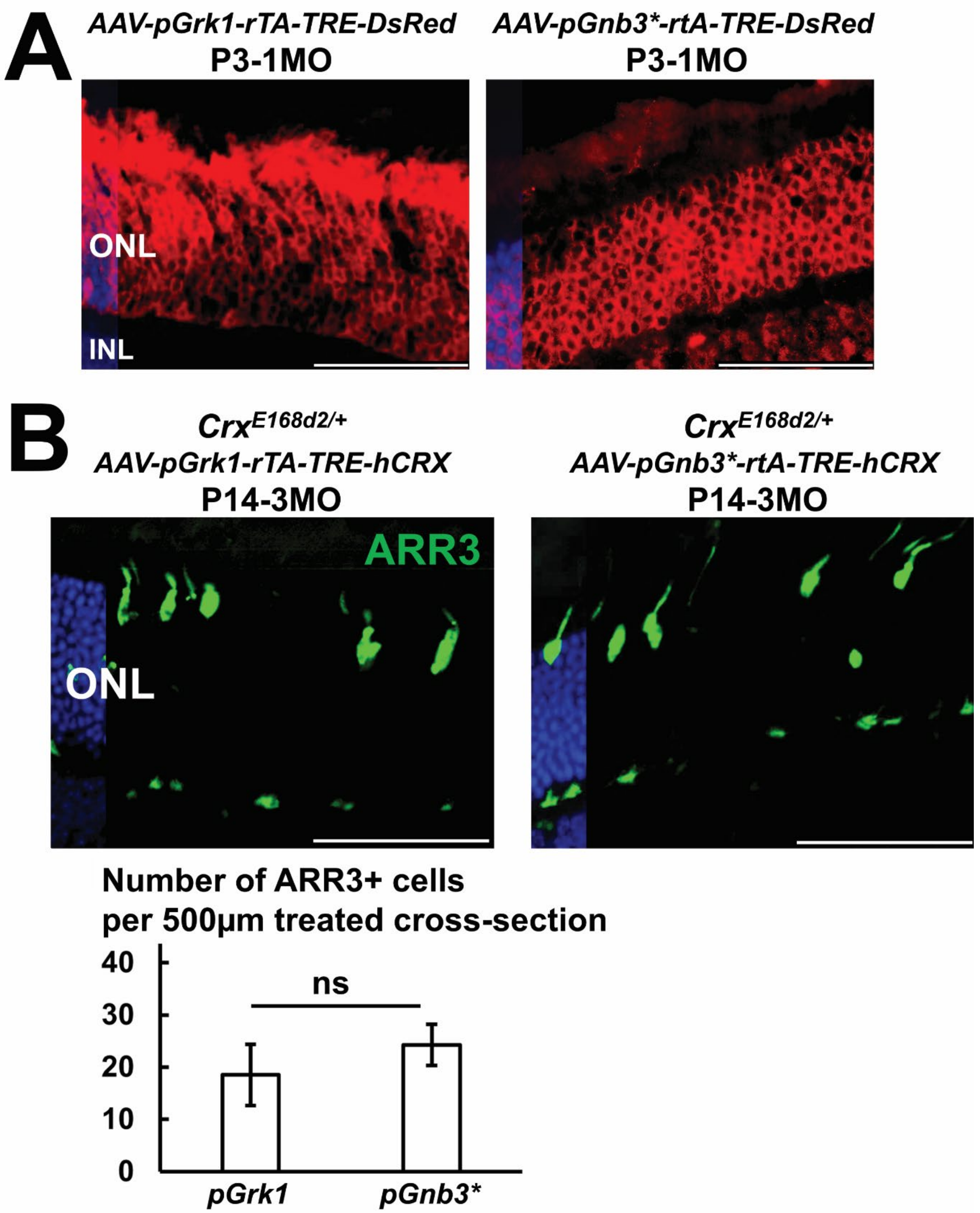

Supplemental figure 13. *AAV* constructs driven by *pGrk1* and *pGnb3**. (A) DsRed-labelled cells in transduced *WT* retinae. Nuclei are stained by DAPI (in blue). Scale bar = 50μm. (B) Anti-ARR3 IHC staining (in green) in $Crx^{E168d2/+;AAV\text{-}hCRX}$ retinae treated by two *AAV* constructs at 3MO. Cell count is based on anti-ARR3 IHC staining within transduced regions. Error bars represent SD on mean values (n=4). Statistical analysis by pairwise t-test. ns means not significant.

Supplemental figure 14

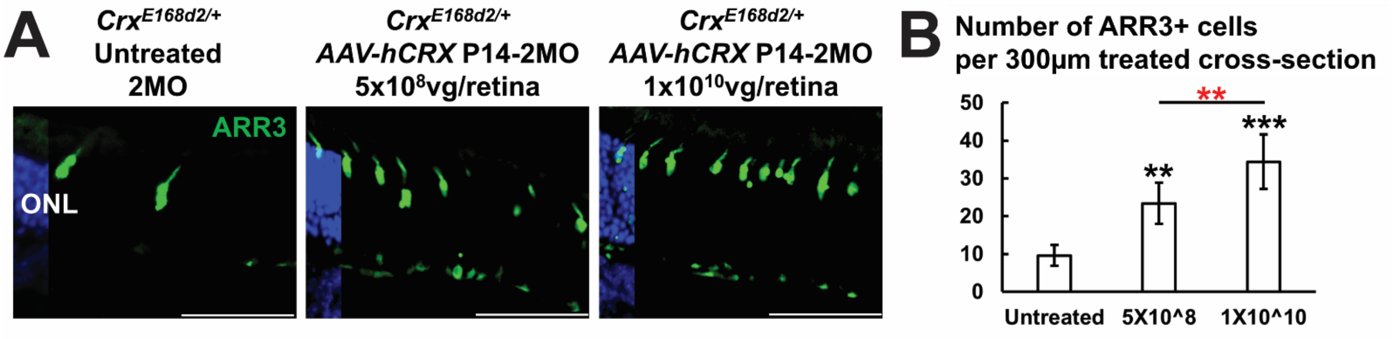

Supplemental figure 14. Testing titers with *AAV-hCRX*. (A) Anti-ARR3 IHC staining (in green) in untreated retinae and $Crx^{E168d2/+;AAV\text{-}hCRX}$ retinae treated by two titers of *pGnb3*-rTA-TRE-hCRX* at 2MO. Nuclei are stained by DAPI (in blue). Scale bar = 50μm. (B) Cone numbers in untreated retinae and $Crx^{E168d2/+;AAV\text{-}hCRX}$ retinae treated by two titers. Cell count is based on anti-ARR3 IHC staining within transduced regions. Error bars represent SD on mean values (n=4). Statistical analysis is performed with ANOVA. Statistical significance indicated by black asterisks compares with untreated retinae with $Crx^{E168d2/+;AAV\text{-}hCRX}$ retinae. Statistical significance indicated by red asterisks compares between retinae treated by two titers. Statistical significance in this figure is indicated by asterisks (**, ***) denoting p≤0.01 and p≤0.001 respectively.

Supplemental figure 15

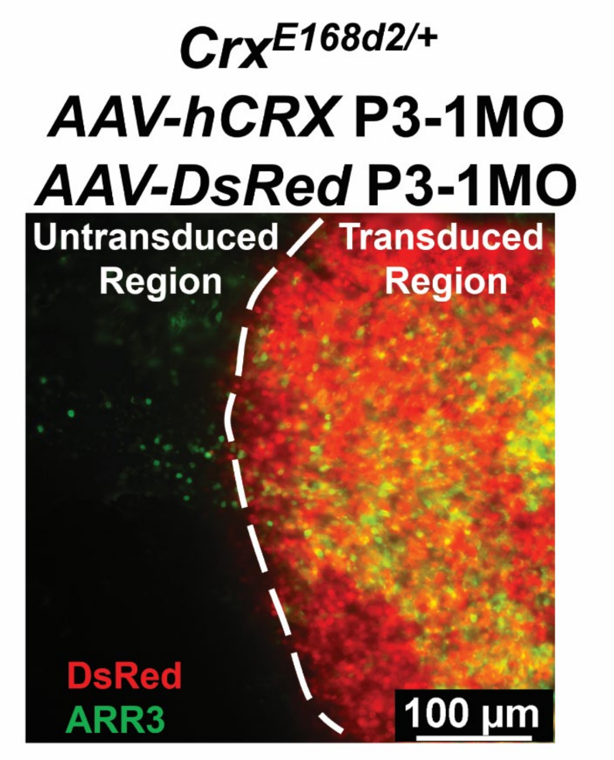

Supplemental figure 15. Whole amount of *Crx*$^{E168d2/+;AAV\text{-}hCRX}$ retinae. Retinae are co-transduced with *pGnb3**-rTA-TRE-hCRX* and *pGnb3*-rTA-TRE-DsRed*. Anti-ARR3 IHC staining is labelled in green. Scale bar = 100μm.

Supplemental figure 16

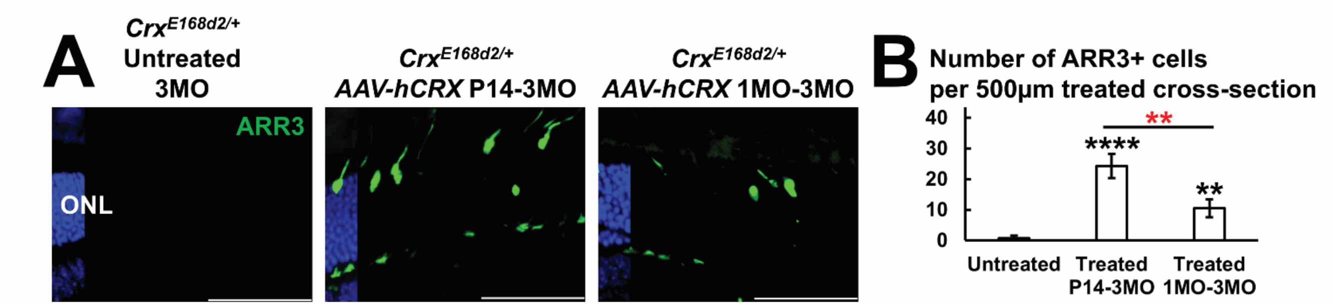

Supplemental figure 16. Early vs late treatment by *AAV-hCRX*-mediated *CRX* augmentation in $Crx^{E168d2/+}$ retinae. Early treatment means P14-3MO, late treatment means 1MO-3MO. (A) Anti-ARR3 IHC staining (in green) of samples treated by different regimes at 3MO. Nuclei are stained by DAPI (in blue). Scale bars = 50µm. (B) Cone numbers in samples treated by different regimes at 3MO. Cell count is based on anti-ARR3 IHC staining within transduced regions. Error bars represent SD on mean values (n=4). Statistical analysis is performed with ANOVA. Statistical significance indicated by black asterisks compares with untreated retinae with $Crx^{E168d2/+;AAV\text{-}hCRX}$ retinae. Statistical significance indicated by red asterisks compares between early and late treatments. Statistical significance in this figure is indicated by asterisks (**, ****) denoting p≤0.01 and p≤0.0001 respectively.

Supplemental figure 17

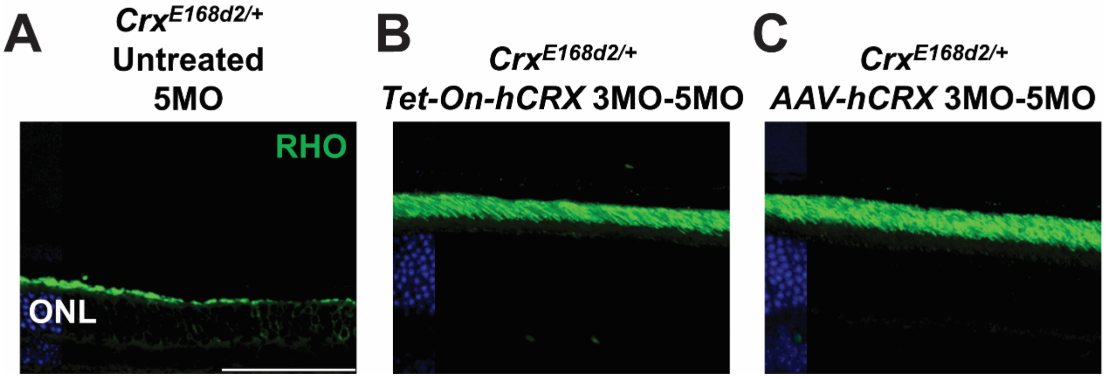

Supplemental figure 17. *CRX* augmentation rescuing late-stage rod degeneration. Treatments begin at 3MO until sample harvest at 5MO. (A) Anti-RHO IHC staining (in green) in $Crx^{E168d2/+}$ retinae at 5MO. Nuclei are stained by DAPI (in blue). Scale bar = 50μm. (B) Anti-RHO IHC staining (in green) in $Crx^{E168d2/+;Tet\text{-}On\text{-}hCRX}$ retinae at 5MO. (C) Anti-RHO IHC staining (in green) in $Crx^{E168d2/+;AAV\text{-}hCRX}$ retinae at 5MO.

Supplemental figure 18

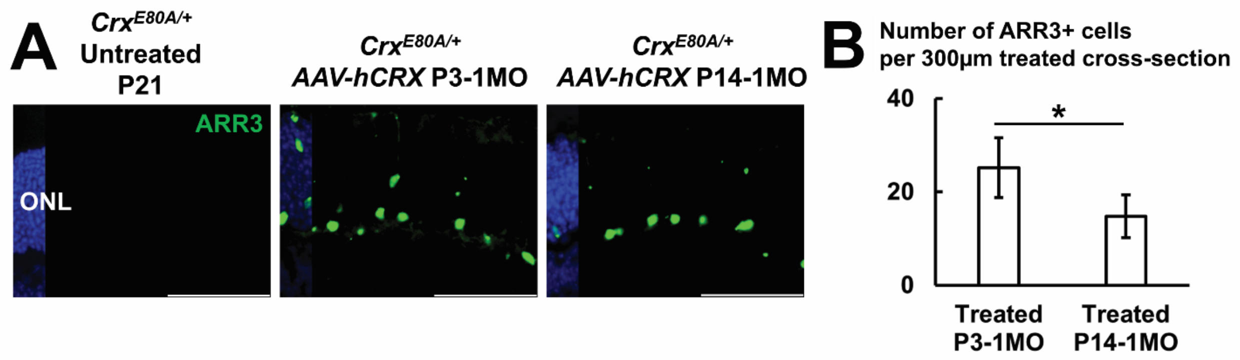

Supplemental figure 18. *AAV-hCRX*-mediated *CRX* augmentation rescuing *Crx*$^{E80A/+}$ retinae. Early treatment means P3-1MO, late treatment means P14-1MO (A) Anti-ARR3 IHC staining (in green) in untreated retinae at P21 and treated by different regimes at 1MO. Nuclei are stained by DAPI (in blue). Scale bars = 50µm. (B) Cone numbers in samples treated by different regimes at 1MO. Cell count is based on anti-ARR3 IHC staining within transduced regions. Error bars represent SD on mean values (n=4). Statistical analysis by pairwise t-test. Statistical significance in this figure is indicated by asterisk (*) denoting p≤0.05.

Supplemental figure 19

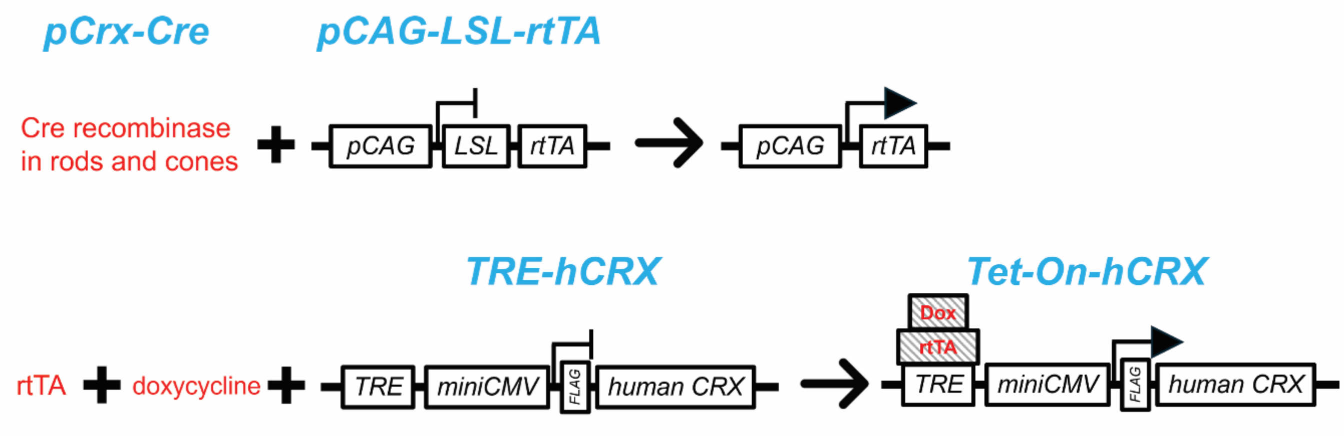

Supplemental figure 19. Schematic diagram of the *Tet-On-hCRX* system.

Supplemental figure 20

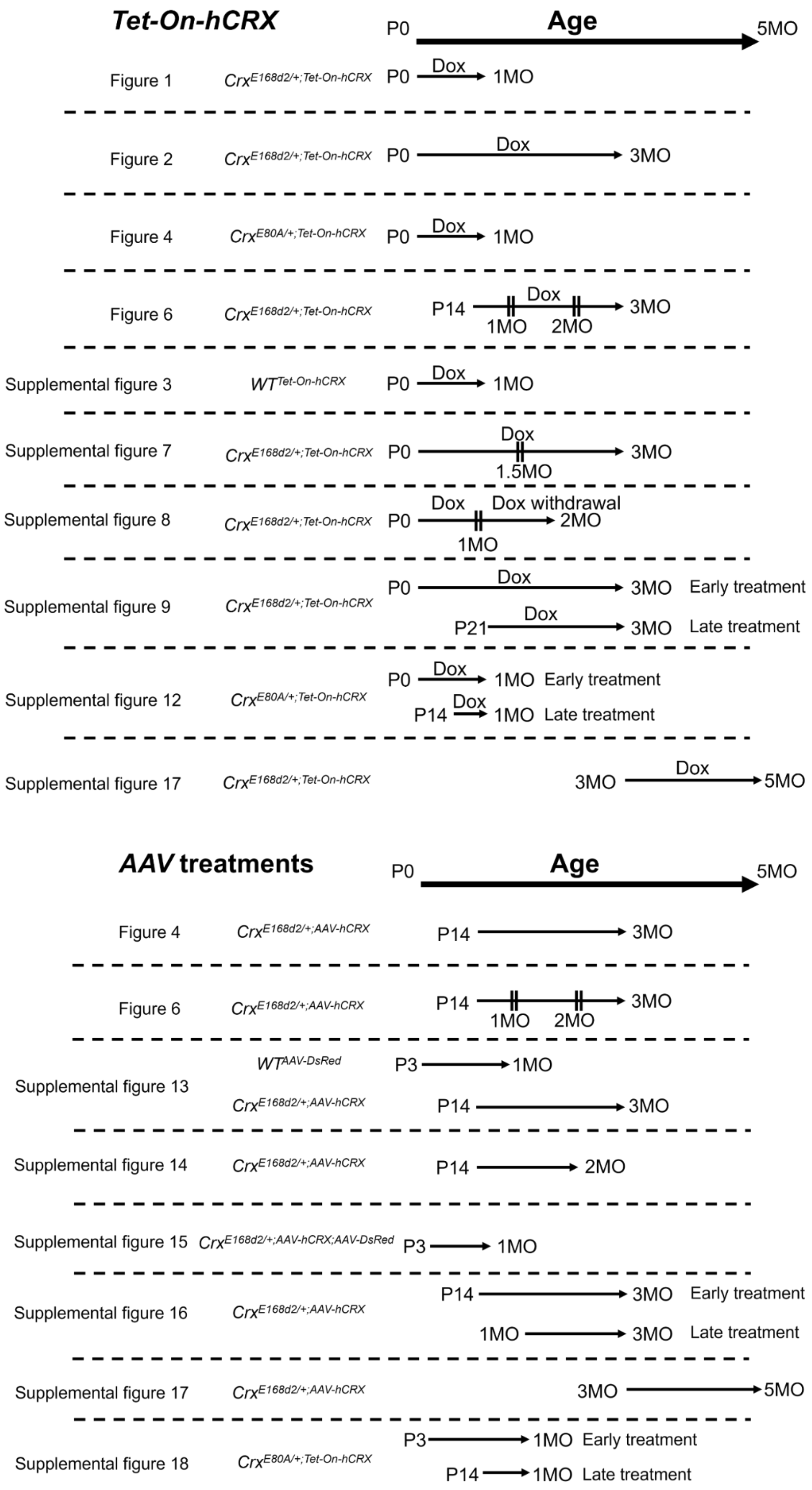

Supplemental figure 20. Treatment schemes in this study.

Supplemental figure 21

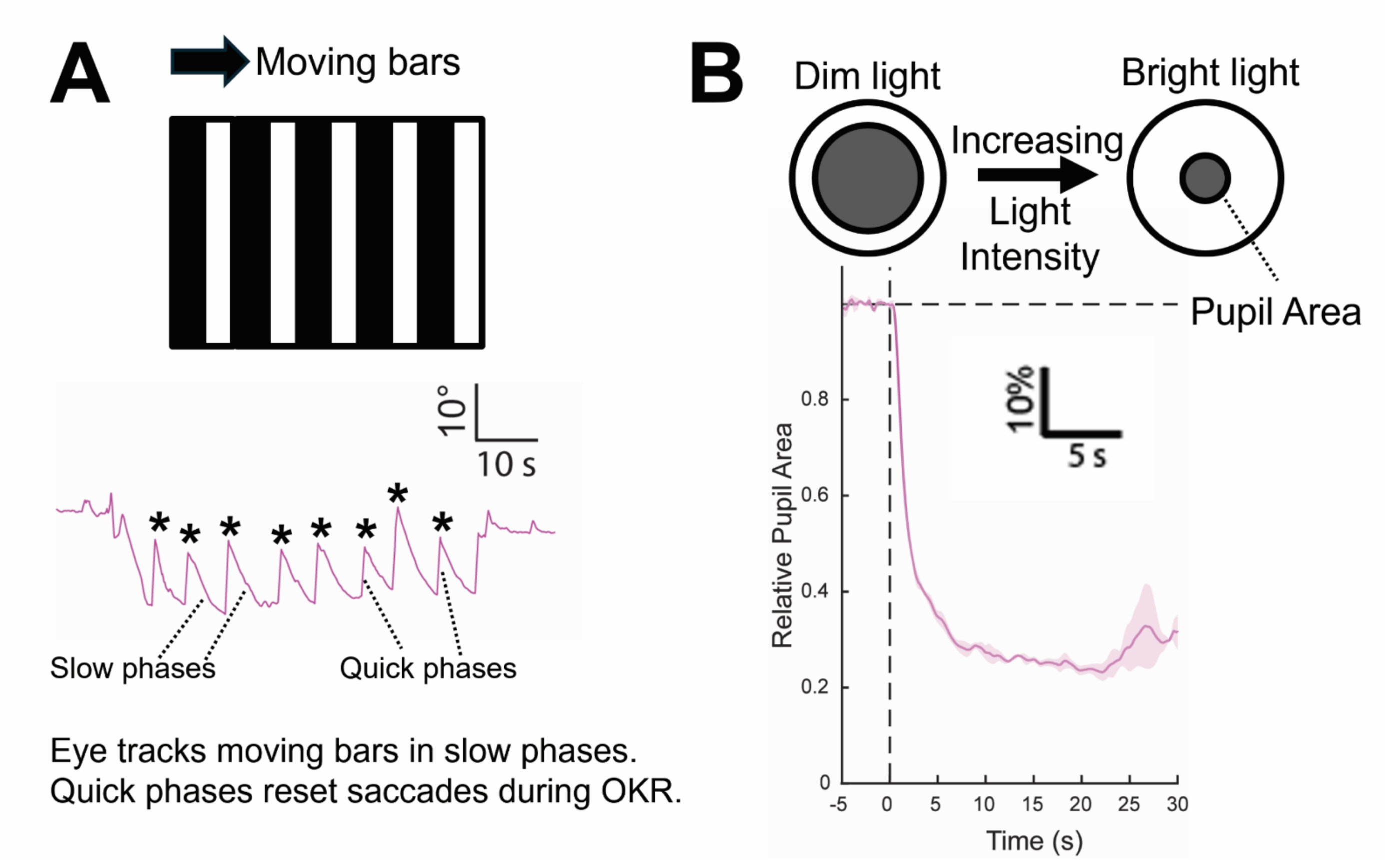

Supplemental figure 21. Introduction of OKR and PLR. (A) OKR measures eye-tracking movements on moving bars. An example of OKR illustrates slow and quick phases as well as numbers of eye-tracking movements (marked by asterisks). (B) PLR records changes in reactive pupil areas in response to altering light intensities.
