## Supplemental Figures and Tables for "Preclinical *CRX* augmentation therapies for *CRX*-associated autosomal dominant cone-rod dystrophies": CRX Augmentation MS Supplemental Table.pdf

Supplemental table 1. *Tet-On-hCRX* sequence

| ***Tet-On-hCRX* (5' to 3')** | |
|---|---|
| GGATCAGGGCAGTCTGGTACTTCCAAGCTCATTAGATGCGTTTAAATAGGAGATCATGCATCCAGATCTGGTGATAG<br>GTGGCAAGTGGTATTCCGTAAGGATATCGTTTTCGGGAGTAGTGCCCCAACTGGGGTAACCTTTGAGTTCTCTCAG<br>TTGGGGGCGTAGGGTCGCCGACGTTTTCGGGAGTAGTGCCCCAACTGGGGTAACCTTTGGGCTCCCCGGGCGCG<br>TACTCCACCTCACCCATCTGGTCCATCATGATGAACGGGTCGAGGTGGCGGTAGTTGATCCCGGCGAACGCGCGG<br>CGCACCGGGAAGCCCTCGCCCTCGAAACCGCTGGGCGCGGTGGTCACGGTGAGCACGGGACGTGCGACGGCGT<br>CGGCGGGTGCGGATACGCGGGGCAGCGTCAGCGGGTTCTCGACGGTCACGGCGGGCATGGGGCCCTGCAGGGG<br>ATCCACCGGTATCGATTTAATTAAGCTTACGCGTCTGCAGCTCGAGGCATGCGTCGACGTTTAAACCATATGGATAT<br>CGTTAACGAATTCATTTAAATTGGGCGCGCCATTTAAATGATATTCGAGTTTACTCCCTATCAGTGATAGAGAACGTA<br>TGTCGAGTTTACTCCCTATCAGTGATAGAGAACGATGTCGAGTTTACTCCCTATCAGTGATAGAGAACGTATGTCGA<br>GTTTACTCCCTATCAGTGATAGAGAACGTATGTCGAGTTTACTCCCTATCAGTGATAGAGAACGTATGTCGAGTTTAT<br>CCCTATCAGTGATAGAGAACGTATGTCGAGTTTACTCCCTATCAGTGATAGAGAACGTATGTCGAGGTAGGCGTGTA<br>CGGTGGGAGGCCTATATAAGCAGAGCTCGTTTAGTGAACCGTCAGATCGCCTGGAGAATTCACGCGTGGTACCTCT<br>AGAGTCGACCCAAGTTGGTCGTGAGGCACTGGGCAGGTAAGTATCAAGGTTACAAGACAGGTTTAAGGAGACCAAT<br>AGAAACTGGGCTTGTCGAGACAGAGAAGACTCTTGCGTTTCTGATAGGCACCTATTGGTCTTACTGACATCCACTTT<br>GCCTTTCTCTCCACAGGTGTCCACTCCCAGTTCAATTACAGCTCTTAAGGCTAGAGTACTTAATACGACTCACTATA<br>GGCTAGCCTCGAGAATTCACGCGTGGTACCTCTAGAGTCGACCCGGGCGGCCGCACCATGGACTACAAAGACCAT<br>GACGGTGATTATAAAGATCATGACATCGATTACAAGGATGACGATGACAAGCTTGCGGCAGCGAATTCCATGGCGT<br>ATATGAACCCAGGGCCCCACTATTCTGTCAACGCCTTGGCCCTAAGTGGCCCCAGTGTGGATCTGATGCACCAGGC<br>TGTGCCCTACCCAAGCGCCCCCAGGAAGCAGCGGCGGGAGCGCACCACCTTCACCCGGAGCCAACTGGAGGAGC<br>TGGAGGCACTGTTTGCCAAGACCCAGTACCCAGACGTCTATGCCCGTGAGGAGGTGGCTCTGAAGATCAATCTGC<br>CTGAGTCCAGGGTTCAGGTTTGGTTCAAGAACCGGAGGGCTAAATGCAGGCAGCAGCGACAGCAGCAGAAACAGC<br>AGCAGCAGCCCCCAGGGGGCCAGGCCAAGGCCCGGCCTGCCAAGAGGAAGGCGGGCACGTCCCCAAGACCCTC<br>CACAGATGTGTGTCCAGACCCTCTGGGCATCTCAGATTCCTACAGTCCCCCTCTGCCCGGCCCCTCAGGCTCCCC<br>AACCACGGCAGTGGCCACTGTGTCCATCTGGAGCCCAGCCTCAGAGTCCCCTTTGCCTGAGGCGCAGCGGGCTG<br>GGCTGGTGGCCTCAGGGCCGTCTCTGACCTCCGCCCCCTATGCCATGACCTACGCCCCGGCCTCCGCTTTCTGCT<br>CTTCCCCCTCCGCCTATGGGTCTCCGAGCTCCTATTTCAGCGGCCTAGACCCCTACCTTTCTCCCATGGTGCCCCA<br>GCTAGGGGGCCCGGCTCTTAGCCCCCTCTCTGGCCCCTCCGTGGGACCTTCCCTGGCCCAGTCCCCCACCTCCCT<br>ATCAGGCCAGAGCTATGGCGCCTACAGCCCCGTGGATAGCTTGGAATTCAAGGACCCCACGGGCACCTGGAAATT<br>CACCTACAATCCCATGGACCCTCTGGACTACAAGGATCAGAGTGCCTGGAAGTTTCAGATCTTGTAGGCGGCCGCT<br>TCGAGCAGACATGATAAGATACATTGATGAGTTTGGACAAACCACAACTAGAATGCAGTGAAAAAAATGCTTTATTTG<br>TGAAATTTGTGATGCTATTGCTTTATTTGTAACCATTATAAGCTGCAATAAACAAGTTAACAACAACAATTGCATTCAT<br>TTTATGTTTCAGGTTCAGGGGGAGATGTGGGAGGTTTTTTAAAGCAAGTAAAACCTCTACAAATGTGGTAAAATCGA<br>TTTAAATGTTTAAACGGCGCGCCCAACTAGTTCTAGAGCGGCCGCCACTCGACGATGTAGGTCACGGTCTCGAAGC<br>CGCGGTGCGGGTGCCAGGGCGTGCCCTTGAGTTCTCTCAGTTGGGGGCGTAGGGTCGCCGACGGATCCTTTGTC<br>GACGAAGTTCCTATTCCGAAGTTCCTATTCTTCAAAAGGTATAGGAACTTCGCGGCCG | ***attP***<br>***TRE***<br>***miniCMV***<br>***Kozak sequence***<br>***FLAG***<br>**Human *CRX* cDNA**<br>***SV40*** |

Supplemental table 2. *AAV* constructs

| ***pGnb3*-Tet-Off-DsRed*** (5' to 3') | |
|---|---|
| AAGACCCTGGGAGTCATGTGTGGATGCATGAGGACAGACCCCAGCCGAGAAGCAGGACACAGAGCTCTGTTCTCT<br>TTTGTAGTCATTTCTTTGCCCACAACACAGGGGCACATGGGCCCAGCTCCCCCCACCCCCATCTCCCTGTCAACTC<br>CCCCAACTATAATTAGGCCCAAGGGCAGCTGGTCGGGGCTTGTCGAGAAAAGGGATTATCCTGCTCAGCCCATCTA<br>ATTACAGCTTCTGCTTGGACCCTCCCCACCCCTCCCCACAGCCCTCAACGAGGAAGCATGGTTTGTAGTGTTGACA<br>ACAGGGTCATCTGTCACCCTCCTGTTCTGGCTCCTGGCCCGCTTGCTGAACAAGGCCAAGATCATGACCACAGGTT<br>GGGGCTAGGGCCTGGGGAGGTCACTGGAGCTATGACTGCTCCTGTTCTACTTCCCCAAACGCAACCAGTAATCCTC<br>ACGGCTTGTCTCTGATCCGTGGGCTGATGCTCTTATCTCTGCTGGCGGACTGACCGTGGAAGCACCCCTCTATCCC<br>CACCCCCCTTCTTGTGGCTGCATAAGAGGGGTGCTGCTCTTCTCTGGTCCACGGGCCTGTTTCCCACCCCTCCAGG<br>CTGACCTGTCTCTGGAAGCCACATACCTCCCCTCCCTGGCCTCTCCCCCACCCCTGCCAGCTGATTTCATTTGCCT<br>GGCGTGGTTGTTGCTGGCCTTACCCTTCCCTGTCATTCCCCCCTCCCTCTCAGTCAGGGCCAGGCCAGGCCAGCT<br>CCTCTGGCAGCAGAGGGGGGCAGGTGACAGGCAGGCATCGCAGCTGAGACAGTGAGGAGGCGGGTACCGCGGG<br>CCCGGGATCCACCGGTTGCCACCATGGCTAGATTAGATAAAAGTAAAGTGATTAACAGCGCATTAGAGCTGCTTAAT<br>GAGGTCGGAATCGAAGGTTTAACAACCCGTAAACTCGCCCAGAAGCTAGGTGTAGAGCAGCCTACATTGTATTGGC<br>ATGTAAAAAATAAGCGGGCTTTGCTCGACGCCTTAGCCATTGAGATGTTAGATAGGCACCATACTCACTTTTGCCCT<br>TTAGAAGGGGAAAGCTGGCAAGATTTTTTACGTAATAACGCTAAAAGTTTTAGATGTGCTTTACTAAGTCATCGCGAT<br>GGAGCAAAAGTACATTTAGGTACACGGCCTACAGAAAAACAGTATGAAACTCTCGAAAATCAATTAGCCTTTTTATG<br>CCAACAAGGTTTTTCACTAGAGAATGCATTATATGCACTCAGCGCTGTGGGGCATTTTACTTTAGGTTGCGTATTGG<br>AAGATCAAGAGCATCAAGTCGCTAAAGAAGAAAGGGAAACACCTACTACTGATAGTATGCCGCCATTATTACGACAA<br>GCTATCGAATTATTTGATCACCAAGGTGCAGAGCCAGCCTTCTTATTCGGCCTTGAATTGATCATATGCGGATTAGA<br>AAAACAACTTAAATGTGAAAGTGGGTCCGCGTACAGCCGCGCGCGTACGAAAAACAATTACGGGTCTACCATCGAG<br>GGCCTGCTCGATCTCCCGGACGACGACGCCCCCGAAGAGGCGGGGCTGGCGGCTCCGCGCCTGTCCTTTCTCCC<br>CGCGGGACACACGCGCAGACTGTCGACGGCCCCCCCGACCGATGTCAGCCTGGGGGACGAGCTCCACTTAGACG<br>GCGAGGACGTGGCGATGGCGCATGCCGACGCGCTAGACGATTTCGATCTGGACATGTTGGGGGACGGGGATTCC<br>CCGGGTCCGGGATTTACCCCCCACGACTCCGCCCCCTACGGCGCTCTGGATATGGCCGACTTCGAGTTTGAGCAG<br>ATGTTTACCGATGCCCTTGGAATTGACGAGTACGGTGGGTAGGGCGCGCCATTTAAATGATATTCGAGTTTACTCCC<br>TATCAGTGATAGAGAACGTATGTCGAGTTTACTCCCTATCAGTGATAGAGAACGATGTCGAGTTTACTCCCTATCAG<br>TGATAGAGAACGTATGTCGAGTTTACTCCCTATCAGTGATAGAGAACGTATGTCGAGTTTACTCCCTATCAGTGATA<br>GAGAACGTATGTCGAGTTTATCCCTATCAGTGATAGAGAACGTATGTCGAGTTTACTCCCTATCAGTGATAGAGAAC<br>GTATGTCGAGGTAGGCGTGTACGGTGGGAGGCCTATATAAGCAGAGCTCGTTTAGTGAACCGTCAGATCGCCTGG<br>AGAATTCACGCGTGGTACCTCTAGAGTCGACCCAAGTTGGTCGTGAGGCACTGGGCAGGTAAGTATCAAGGTTACA<br>AGACAGGTTTAAGGAGACCAATAGAAACTGGGCTTGTCGAGACAGAGAAGACTCTTGCGTTTCTGATAGGCACCTA<br>TTGGTCTTACTGACATCCACTTTGCCTTTCTCTCCACAGGTGTCCACTCCCAGTTCAATTACAGCTCTTAAGGCTCC<br>GGTCGCCACCATGGCCTCCTCCGAGAACGTCATCACCGAGTTCATGCGCTTCAAGGTGCGCATGGAGGGCACCGT<br>GAACGGCCACGAGTTCGAGATCGAGGGCGAGGGCGAGGGCCGCCCCTACGAGGGCCACAACACCGTGAAGCTGA<br>AGGTGACCAAGGGCGGCCCCCTGCCCTTCGCCTGGGACATCCTGTCCCCCCAGTTCCAGTACGGCTCCAAGGTGT<br>ACGTGAAGCACCCCGCCGACATCCCCGACTACAAGAAGCTGTCCTTCCCCGAGGGCTTCAAGTGGGAGCGCGTGA<br>TGAACTTCGAGGACGGCGGCGTGGCGACCGTGACCCAGGACTCCTCCCTGCAGGACGGCTGCTTCATCTACAAGG<br>TGAAGTTCATCGGCGTGAACTTCCCCTCCGACGGCCCCGTGATGCAGAAGAAGACCATGGGCTGGGAGGCCTCCA<br>CCGAGCGCCTGTACCCCCGCGACGGCGTGCTGAAGGGCGAGACCCACAAGGCCCTGAAGCTGAAGGACGGCGG<br>CCACTACCTGGTGGAGTTCAAGTCCATCTACATGGCCAAGAAGCCCGTGCAGCTGCCCGGCTACTACTACGTGGA<br>CGCCAAGCTGGACATCACCTCCCACAACGAGGACTACACCATCGTGGAGCAGTACGAGCGCACCGAGGGCCGCC<br>ACCACCTGTTCCTGTAGCGGCCGCGACTCTAGATCATAATCAGCCATACCACATTTGTAGAGGTTTTACTTGCTTTA<br>AAAAACCTCCCACACCTCCCCCTGAACCTGAAACATAAAATGAATGCAATTGTTGTTGT | ***pGnb3****<br>***Kozak sequence***<br>***rTA***<br>***TRE***<br>***FLAG***<br>***DsRed*** |

### *pGrk1-Tet-Off-DsRed* (5' to 3')

GGGCCCCAGAAGCCTGGTGGTTGTTTGTCCTTCTCAGGGGAAAAGTGAGGCGGCCCCTTGGAGGAAGGGGCCGG
GCAGAATGATCTAATCGGATTCCAAGCAGCTCAGGGGATTGTCTTTTTCTAGCACCTTCTTGCCACTCCTAAGCGTC
CTCCGTGACCCCGGCTGGGATTTAGCCTGGTGCTGTGTCAGCCCCGGGCTCCCAGGGGCTTCCCAGTGGTCCCCA
GGAACCCTCGACAGGGCCAGGGCGTCTCTCTCGTCCAGCAAGGGCAGGGACGGGCCACAGGCCAAGGGCGGTAC
CGCGGGCCCGGGATCCACCGGTGCCACCATGGCTAGATTAGATAAAAGTAAAGTGATTAACAGCGCATTAGAGCTG
CTTAATGAGGTCGGAATCGAAGGTTTAACAACCCGTAAACTCGCCCAGAAGCTAGGTGTAGAGCAGCCTACATTGT
ATTGGCATGTAAAAAATAAGCGGGCTTTGCTCGACGCCTTAGCCATTGAGATGTTAGATAGGCACCATACTCACTTT
TGCCCTTTAGAAGGGGAAAGCTGGCAAGATTTTTTACGTAATAACGCTAAAAGTTTTAGATGTGCTTTACTAAGTCAT
CGCGATGGAGCAAAAGTACATTTAGGTACACGGCCTACAGAAAAACAGTATGAAACTCTCGAAAATCAATTAGCCTT
TTTATGCCAACAAGGTTTTTCACTAGAGAATGCATTATATGCACTCAGCGCTGTGGGGCATTTTACTTTAGGTTGCGT
ATTGGAAGATCAAGAGCATCAAGTCGCTAAAGAAGAAAGGGAAACACCTACTACTGATAGTATGCCGCCATTATTAC
GACAAGCTATCGAATTATTTGATCACCAAGGTGCAGAGCCAGCCTTCTTATTCGGCCTTGAATTGATCATATGCGGA
TTAGAAAAACAACTTAAATGTGAAAGTGGGTCCGCGTACAGCCGCGCGCGTACGAAAAACAATTACGGGTCTACCA
TCGAGGGCCTGCTCGATCTCCCGGACGACGACGCCCCCGAAGAGGCGGGGCTGGCGGCTCCGCGCCTGTCCTTT
CTCCCCGCGGGACACACGCGCAGACTGTCGACGGCCCCCCCGACCGATGTCAGCCTGGGGGACGAGCTCCACTT
AGACGGCGAGGACGTGGCGATGGCGCATGCCGACGCGCTAGACGATTTCGATCTGGACATGTTGGGGGACGGGG
ATTCCCCGGGTCCGGGATTTACCCCCCACGACTCCGCCCCCTACGGCGCTCTGGATATGGCCGACTTCGAGTTTG
AGCAGATGTTTACCGATGCCCTTGGAATTGACGAGTACGGTGGGTAGGGCGCGCCATTTAAATGATATTCGAGTTT
ACTCCCTATCAGTGATAGAGAACGTATGTCGAGTTTACTCCCTATCAGTGATAGAGAACGATGTCGAGTTTACTCCC
TATCAGTGATAGAGAACGTATGTCGAGTTTACTCCCTATCAGTGATAGAGAACGTATGTCGAGTTTACTCCCTATCA
GTGATAGAGAACGTATGTCGAGTTTATCCCTATCAGTGATAGAGAACGTATGTCGAGTTTACTCCCTATCAGTGATA
GAGAACGTATGTCGAGGTAGGCGTGTACGGTGGGAGGCCTATATAAGCAGAGCTCGTTTAGTGAACCGTCAGATC
GCCTGGAGAATTCACGCGTGGTACCTCTAGAGTCGACCCAAGTTGGTCGTGAGGCACTGGGCAGGTAAGTATCAA
GGTTACAAGACAGGTTTAAGGAGACCAATAGAAACTGGGCTTGTCGAGACAGAGAAGACTCTTGCGTTTCTGATAG
GCACCTATTGGTCTTACTGACATCCACTTTGCCTTTCTCTCCACAGGTGTCCACTCCCAGTTCAATTACAGCTCTTAA
GGCTCCGGTCGCCACCATGGCCTCCTCCGAGAACGTCATCACCGAGTTCATGCGCTTCAAGGTGCGCATGGAGGG
CACCGTGAACGGCCACGAGTTCGAGATCGAGGGCGAGGGCGAGGGCCGCCCCTACGAGGGCCACAACACCGTGA
AGCTGAAGGTGACCAAGGGCGGCCCCCTGCCCTTCGCCTGGGACATCCTGTCCCCCCAGTTCCAGTACGGCTCCA
AGGTGTACGTGAAGCACCCCGCCGACATCCCCGACTACAAGAAGCTGTCCTTCCCCGAGGGCTTCAAGTGGGAGC
GCGTGATGAACTTCGAGGACGGCGGCGTGGCGACCGTGACCCAGGACTCCTCCCTGCAGGACGGCTGCTTCATCT
ACAAGGTGAAGTTCATCGGCGTGAACTTCCCCTCCGACGGCCCCGTGATGCAGAAGAAGACCATGGGCTGGGAGG
CCTCCACCGAGCGCCTGTACCCCCGCGACGGCGTGCTGAAGGGCGAGACCCACAAGGCCCTGAAGCTGAAGGAC
GGCGGCCACTACCTGGTGGAGTTCAAGTCCATCTACATGGCCAAGAAGCCCGTGCAGCTGCCCGGCTACTACTAC
GTGGACGCCAAGCTGGACATCACCTCCCACAACGAGGACTACACCATCGTGGAGCAGTACGAGCGCACCGAGGG
CCGCCACCACCTGTTCCTGTAGCGGCCGCGACTCTAGATCATAATCAGCCATACCACATTTGTAGAGGTTTTACTTG
CTTTAAAAAACCTCCCACACCTCCCCCTGAACCTGAAACATAAAATGAATGCAATTGTTGTTGT

***pGrk1***
***Kozak sequence***
***rTA***
***TRE***
***FLAG***
***DsRed***

### *pGnb3*-Tet-Off-hCRX* (5' to 3')

AAGACCCTGGGAGTCATGTGTGGATGCATGAGGACAGACCCCAGCCGAGAAGCAGGACACAGAGCTCTGTTCTCT
TTTGTAGTCATTTCTTTGCCCACAACACAGGGGCACATGGGCCCAGCTCCCCCCACCCCCATCTCCCTGTCAACTC
CCCCAACTATAATTAGGCCCAAGGGCAGCTGGTCGGGGCTTGTCGAGAAAAGGGATTATCCTGCTCAGCCCATCTA
ATTACAGCTTCTGCTTGGACCCTCCCCACCCCTCCCCACAGCCCTCAACGAGGAAGCATGGTTTGTAGTGTTGACA
ACAGGGTCATCTGTCACCCTCCTGTTCTGGCTCCTGGCCCGCTTGCTGAACAAGGCCAAGATCATGACCACAGGTT
GGGGCTAGGGCCTGGGGAGGTCACTGGAGCTATGACTGCTCCTGTTCTACTTCCCCAAACGCAACCAGTAATCCTC
ACGGCTTGTCTCTGATCCGTGGGCTGATGCTCTTATCTCTGCTGGCGGACTGACCGTGGAAGCACCCCTCTATCCC
CACCCCCCTTCTTGTGGCTGCATAAGAGGGGTGCTGCTCTTCTCTGGTCCACGGGCCTGTTTCCCACCCCTCCAGG
CTGACCTGTCTCTGGAAGCCACATACCTCCCCTCCCTGGCCTCTCCCCCACCCCTGCCAGCTGATTTCATTTGCCT
GGCGTGGTTGTTGCTGGCCTTACCCTTCCCTGTCATTCCCCCCTCCCTCTCAGTCAGGGCCAGGCCAGGCCAGCT
CCTCTGGCAGCAGAGGGGGGCAGGTGACAGGCAGGCATCGCAGCTGAGACAGTGAGGAGGCGGGTACCGCGGG
CCCGGGATCCACCGGTTGCCACCATGGCTAGATTAGATAAAAGTAAAGTGATTAACAGCGCATTAGAGCTGCTTAAT
GAGGTCGGAATCGAAGGTTTAACAACCCGTAAACTCGCCCAGAAGCTAGGTGTAGAGCAGCCTACATTGTATTGGC
ATGTAAAAAATAAGCGGGCTTTGCTCGACGCCTTAGCCATTGAGATGTTAGATAGGCACCATACTCACTTTTGCCCT
TTAGAAGGGGAAAGCTGGCAAGATTTTTTACGTAATAACGCTAAAAGTTTTAGATGTGCTTTACTAAGTCATCGCGAT
GGAGCAAAAGTACATTTAGGTACACGGCCTACAGAAAAACAGTATGAAACTCTCGAAAATCAATTAGCCTTTTTATG
CCAACAAGGTTTTTCACTAGAGAATGCATTATATGCACTCAGCGCTGTGGGGCATTTTACTTTAGGTTGCGTATTGG
AAGATCAAGAGCATCAAGTCGCTAAAGAAGAAAGGGAAACACCTACTACTGATAGTATGCCGCCATTATTACGACAA
GCTATCGAATTATTTGATCACCAAGGTGCAGAGCCAGCCTTCTTATTCGGCCTTGAATTGATCATATGCGGATTAGA
AAAACAACTTAAATGTGAAAGTGGGTCCGCGTACAGCCGCGCGCGTACGAAAAACAATTACGGGTCTACCATCGAG
GGCCTGCTCGATCTCCCGGACGACGACGCCCCCGAAGAGGCGGGGCTGGCGGCTCCGCGCCTGTCCTTTCTCCC
CGCGGGACACACGCGCAGACTGTCGACGGCCCCCCCGACCGATGTCAGCCTGGGGGACGAGCTCCACTTAGACG
GCGAGGACGTGGCGATGGCGCATGCCGACGCGCTAGACGATTTCGATCTGGACATGTTGGGGGACGGGGATTCC
CCGGGTCCGGGATTTACCCCCCACGACTCCGCCCCCTACGGCGCTCTGGATATGGCCGACTTCGAGTTTGAGCAG
ATGTTTACCGATGCCCTTGGAATTGACGAGTACGGTGGGTAGGGCGCGCCATTTAAATGATATTCGAGTTTACTCCC
TATCAGTGATAGAGAACGTATGTCGAGTTTACTCCCTATCAGTGATAGAGAACGATGTCGAGTTTACTCCCTATCAG
TGATAGAGAACGTATGTCGAGTTTACTCCCTATCAGTGATAGAGAACGTATGTCGAGTTTACTCCCTATCAGTGATA
GAGAACGTATGTCGAGTTTATCCCTATCAGTGATAGAGAACGTATGTCGAGTTTACTCCCTATCAGTGATAGAGAAC
GTATGTCGAGGTAGGCGTGTACGGTGGGAGGCCTATATAAGCAGAGCTCGTTTAGTGAACCGTCAGATCGCCTGG
AGAATTCACGCGTGGTACCTCTAGAGTCGACCCAAGTTGGTCGTGAGGCACTGGGCAGGTAAGTATCAAGGTTACA
AGACAGGTTTAAGGAGACCAATAGAAACTGGGCTTGTCGAGACAGAGAAGACTCTTGCGTTTCTGATAGGCACCTA
TTGGTCTTACTGACATCCACTTTGCCTTTCTCTCCACAGGTGTCCACTCCCAGTTCAATTACAGCTCTTAAGGCTAGA
GTACTTAATACGACTCACTATAGGCTAGCCTCGAGAATTCACGCGTGGTACCTCTAGAGTCGACCCGGGCGGCCGC
ACCATGGACTACAAAGACCATGACGGTGATTATAAAGATCATGACATCGATTACAAGGATGACGATGACAAGCTTGC
GGCAGCGAATTCCATGGCGTATATGAACCCAGGGCCCCACTATTCTGTCAACGCCTTGGCCCTAAGTGGCCCCAGT
GTGGATCTGATGCACCAGGCTGTGCCCTACCCAAGCGCCCCCAGGAAGCAGCGGCGGGAGCGCACCACCTTCAC
CCGGAGCCAACTGGAGGAGCTGGAGGCACTGTTTGCCAAGACCCAGTACCCAGACGTCTATGCCCGTGAGGAGGT
GGCTCTGAAGATCAATCTGCCTGAGTCCAGGGTTCAGGTTTGGTTCAAGAACCGGAGGGCTAAATGCAGGCAGCA
GCGACAGCAGCAGAAACAGCAGCAGCAGCCCCCAGGGGGCCAGGCCAAGGCCCGGCCTGCCAAGAGGAAGGCG
GGCACGTCCCCAAGACCCTCCACAGATGTGTGTCCAGACCCTCTGGGCATCTCAGATTCCTACAGTCCCCCTCTGC
CCGGCCCCTCAGGCTCCCCAACCACGGCAGTGGCCACTGTGTCCATCTGGAGCCCAGCCTCAGAGTCCCCTTTGC
CTGAGGCGCAGCGGGCTGGGCTGGTGGCCTCAGGGCCGTCTCTGACCTCCGCCCCCTATGCCATGACCTACGCC
CCGGCCTCCGCTTTCTGCTCTTCCCCCTCCGCCTATGGGTCTCCGAGCTCCTATTTCAGCGGCCTAGACCCCTACC
TTTCTCCCATGGTGCCCCAGCTAGGGGGCCCGGCTCTTAGCCCCCTCTCTGGCCCCTCCGTGGGACCTTCCCTGG
CCCAGTCCCCCACCTCCCTATCAGGCCAGAGCTATGGCGCCTACAGCCCCGTGGATAGCTTGGAATTCAAGGACC
CCACGGGCACCTGGAAATTCACCTACAATCCCATGGACCCTCTGGACTACAAGGATCAGAGTGCCTGGAAGTTTCA
GATCTTGTAGGCGGCCGC

***pGnb3****
***Kozak sequence***
***rTA***
***TRE***
***FLAG***
**Human *CRX* cDNA**

### *pGnb3*-Tet-Off-DsRed* (5’ to 3’)

AAGACCCTGGGAGTCATGTGTGGATGCATGAGGACAGACCCCAGCCGAGAAGCAGGACACAGAGCTCTGTTCTCT
TTTGTAGTCATTTCTTTGCCCACAACACAGGGGCACATGGGCCCAGCTCCCCCCACCCCCATCTCCCTGTCAACTC
CCCCAACTATAATTAGGCCCAAGGGCAGCTGGTCGGGGCTTGTCGAGAAAAGGGATTATCCTGCTCAGCCCATCTA
ATTACAGCTTCTGCTTGGACCCTCCCCACCCCTCCCCACAGCCCTCAACGAGGAAGCATGGTTTGTAGTGTTGACA
ACAGGGTCATCTGTCACCCTCCTGTTCTGGCTCCTGGCCCGCTTGCTGAACAAGGCCAAGATCATGACCACAGGTT
GGGGCTAGGGCCTGGGGAGGTCACTGGAGCTATGACTGCTCCTGTTCTACTTCCCCAAACGCAACCAGTAATCCTC
ACGGCTTGTCTCTGATCCGTGGGCTGATGCTCTTATCTCTGCTGGCGGACTGACCGTGGAAGCACCCCTCTATCCC
CACCCCCCTTCTTGTGGCTGCATAAGAGGGGTGCTGCTCTTCTCTGGTCCACGGGCCTGTTTCCCACCCCTCCAGG
CTGACCTGTCTCTGGAAGCCACATACCTCCCCTCCCTGGCCTCTCCCCCACCCCTGCCAGCTGATTTCATTTGCCT
GGCGTGGTTGTTGCTGGCCTTACCCTTCCCTGTCATTCCCCCCTCCCTCTCAGTCAGGGCCAGGCCAGGCCAGCT
CCTCTGGCAGCAGAGGGGGGCAGGTGACAGGCAGGCATCGCAGCTGAGACAGTGAGGAGGCGGGTACCGCGGG
CCCGGGATCCACCGGTTGCCACCATGGCTAGATTAGATAAAAGTAAAGTGATTAACAGCGCATTAGAGCTGCTTAAT
GAGGTCGGAATCGAAGGTTTAACAACCCGTAAACTCGCCCAGAAGCTAGGTGTAGAGCAGCCTACATTGTATTGGC
ATGTAAAAAATAAGCGGGCTTTGCTCGACGCCTTAGCCATTGAGATGTTAGATAGGCACCATACTCACTTTTGCCCT
TTAGAAGGGGAAAGCTGGCAAGATTTTTTACGTAATAACGCTAAAAGTTTTAGATGTGCTTTACTAAGTCATCGCGAT
GGAGCAAAAGTACATTTAGGTACACGGCCTACAGAAAAACAGTATGAAACTCTCGAAAATCAATTAGCCTTTTTATG
CCAACAAGGTTTTTCACTAGAGAATGCATTATATGCACTCAGCGCTGTGGGGCATTTTACTTTAGGTTGCGTATTGG
AAGATCAAGAGCATCAAGTCGCTAAAGAAGAAAGGGAAACACCTACTACTGATAGTATGCCGCCATTATTACGACAA
GCTATCGAATTATTTGATCACCAAGGTGCAGAGCCAGCCTTCTTATTCGGCCTTGAATTGATCATATGCGGATTAGA
AAAACAACTTAAATGTGAAAGTGGGTCCGCGTACAGCCGCGCGCGTACGAAAAACAATTACGGGTCTACCATCGAG
GGCCTGCTCGATCTCCCGGACGACGACGCCCCCGAAGAGGCGGGGCTGGCGGCTCCGCGCCTGTCCTTTCTCCC
CGCGGGACACACGCGCAGACTGTCGACGGCCCCCCCGACCGATGTCAGCCTGGGGGACGAGCTCCACTTAGACG
GCGAGGACGTGGCGATGGCGCATGCCGACGCGCTAGACGATTTCGATCTGGACATGTTGGGGGACGGGGATTCC
CCGGGTCCGGGATTTACCCCCCACGACTCCGCCCCCTACGGCGCTCTGGATATGGCCGACTTCGAGTTTGAGCAG
ATGTTTACCGATGCCCTTGGAATTGACGAGTACGGTGGGTAGGGCGCGCCATTTAAATGATATTCGAGTTTACTCCC
TATCAGTGATAGAGAACGTATGTCGAGTTTACTCCCTATCAGTGATAGAGAACGATGTCGAGTTTACTCCCTATCAG
TGATAGAGAACGTATGTCGAGTTTACTCCCTATCAGTGATAGAGAACGTATGTCGAGTTTACTCCCTATCAGTGATA
GAGAACGTATGTCGAGTTTATCCCTATCAGTGATAGAGAACGTATGTCGAGTTTACTCCCTATCAGTGATAGAGAAC
GTATGTCGAGGTAGGCGTGTACGGTGGGAGGCCTATATAAGCAGAGCTCGTTTAGTGAACCGTCAGATCGCCTGG
AGAATTCACGCGTGGTACCTCTAGAGTCGACCCAAGTTGGTCGTGAGGCACTGGGCAGGTAAGTATCAAGGTTACA
AGACAGGTTTAAGGAGACCAATAGAAACTGGGCTTGTCGAGACAGAGAAGACTCTTGCGTTTCTGATAGGCACCTA
TTGGTCTTACTGACATCCACTTTGCCTTTCTCTCCACAGGTGTCCACTCCCAGTTCAATTACAGCTCTTAAGGCTCC
GGTCGCCACCATGGCCTCCTCCGAGAACGTCATCACCGAGTTCATGCGCTTCAAGGTGCGCATGGAGGGCACCGT
GAACGGCCACGAGTTCGAGATCGAGGGCGAGGGCGAGGGCCGCCCCTACGAGGGCCACAACACCGTGAAGCTGA
AGGTGACCAAGGGCGGCCCCCTGCCCTTCGCCTGGGACATCCTGTCCCCCCAGTTCCAGTACGGCTCCAAGGTGT
ACGTGAAGCACCCCGCCGACATCCCCGACTACAAGAAGCTGTCCTTCCCCGAGGGCTTCAAGTGGGAGCGCGTGA
TGAACTTCGAGGACGGCGGCGTGGCGACCGTGACCCAGGACTCCTCCCTGCAGGACGGCTGCTTCATCTACAAGG
TGAAGTTCATCGGCGTGAACTTCCCCTCCGACGGCCCCGTGATGCAGAAGAAGACCATGGGCTGGGAGGCCTCCA
CCGAGCGCCTGTACCCCCGCGACGGCGTGCTGAAGGGCGAGACCCACAAGGCCCTGAAGCTGAAGGACGGCGG
CCACTACCTGGTGGAGTTCAAGTCCATCTACATGGCCAAGAAGCCCGTGCAGCTGCCCGGCTACTACTACGTGGA
CGCCAAGCTGGACATCACCTCCCACAACGAGGACTACACCATCGTGGAGCAGTACGAGCGCACCGAGGGCCGCC
ACCACCTGTTCCTGTAGCGGCCGCGACTCTAGATCATAATCAGCCATACCACATTTGTAGAGGTTTTACTTGCTTTA
AAAAACCTCCCACACCTCCCCCTGAACCTGAAACATAAAATGAATGCAATTGTTGTTGT

*pGnb3**
*Kozak sequence*
*rTA*
*TRE*
*FLAG*
*DsRed*

Supplemental table 3. qPCR primers

| qPCR primers | | |
|---|---|---|
| **Gene** | **Forward Primer (5' – 3')** | **Reverse Primer (5' – 3')** |
| *Actb* | CCAACTGGGACGACATGGAG | TGGTACGACCAGAGGCATACA |
| *Crx* | TGTCCCATACTCAAGTGCCC | TGCTGTTTCTGCTGCTGTCG |
| *Gnat1* | ACGATGGACCTAACACTTACGAGG | TGGAAAGGACGGTATTTGAGG |
| *Gnat2* | AGTCAAGACAACAGGCATCATCG | TCACTTCGTCATCCTCCACCAG |
| *Opn1mw* | GGTGGTGATGGTCTTCGCATAC | TTGGAGGTGCTGGAAAGTTCAG |
| *Opn1sw* | GCTGGACTTACGGCTTGTCACC | TGTGGCGTTGTGTTTGCTGC |
| *Rho* | GCTTCCCTACGCCAGTGTG | CAGTGGATTCTTGCCGCAG |
| *Ubb* | CAACATCCAGAAAGAGTCAACC | ATGTTGTAATCAGAGAGGGTGC |

Supplemental table 4. Antibody information

| Antibody | Experiment | Product Information | Host |
|---|---|---|---|
| RXRG | IHC | Santa Cruz Biotechnology, sc-555 | Rabbit polyclonal |
| RHO | IHC | MilliporeSigma, O4886 | Mouse monoclonal |
| ARR3 | IHC | MilliporeSigma, AB15282 | Rabbit polyclonal |
| ARR3 | Western blotting | Proteintech 11100-2-AP | Rabbit polyclonal |
| Cleaved caspase 3 | IHC | Cell Signaling Technology, 9661 | Rabbit polyclonal |
| GFAP | IHC | MilliporeSigma, G3893 | Mouse monoclonal |
| FLAG | IHC & Western blotting | MilliporeSigma, F1804 | Mouse monoclonal |
| Lamin B1 | Western blotting | Abcam, ab16048 | Rabbit polyclonal |
